# Penetrance estimation of *SORL1* loss-of-function variants using a family-based strategy adjusted on *APOE* genotypes suggest a non-monogenic inheritance

**DOI:** 10.1101/2021.06.30.450554

**Authors:** Catherine Schramm, Camille Charbonnier, Aline Zaréa, Morgane Lacour, David Wallon, CNRMAJ collaborators, Anne Boland, Jean-François Deleuze, Robert Olaso, Flora Alarcon, Dominique Campion, Grégory Nuel, Gaël Nicolas

## Abstract

For complex disorders, estimating the age-related penetrance associated with rare variants of strong effect is essential before a putative use for genetic counseling or disease prevention. However, rarity and co-occurrence with other risk factors make such estimations difficult. In the context of Alzheimer disease, we present a survival model to estimate the penetrance of *SORL1* rare (allele frequency< 1%) Loss-of-Function variants (LoF) while accounting for *APOE*-ε4, the main risk factor (allele frequency∼ 14% in Caucasians). We developed an efficient strategy to compute penetrance estimates accounting for both common and rare genetic variants based on available penetrance curves associated with common risk factors and using incomplete pedigree data to quantify the additional risk conferred by rare variants. Our model combines: (i) a baseline for non-carriers of *SORL1* LoF variants, stratified by *APOE* genotypes derived from the Rotterdam study and (ii) an age-dependent proportional hazard effect for *SORL1* LoF variants estimated from pedigrees with a proband carrying such a variant. We embed this model into an Expectation-Maximisation algorithm to accommodate for missing genotypes. Confidence intervals were computed by bootstraps. To correct for ascertainment bias, proband phenotypes were omitted. We obtained penetrance curves associated with *SORL1* LoF variants at the digenic level. By age 70, we estimate a 100% penetrance of *SORL1* LoF variants only among *APOE*-ε4ε4 carriers, while penetrance is 56%[40% − 72%] among ε4 heterozygous carriers and 37%[26% − 51%] among ε4 non-carriers. We conclude that rare *SORL1* LoF variants should not be used for genetic counseling regardless of the *APOE* status.

## Introduction

In the context of complex disorders, large scale case-control studies based on exome and genome sequencing have enabled the discovery of rare risk variants with strong effects. Such variants could not be caught by genome-wide association studies (GWAS) performed on DNA chips^1^. While the variant effects are usually presented as odds ratios or relative risks, absolute penetrance estimates, i.e. the probability of developing the disease according to age for carriers of a given genotype, are much more informative in a clinical setting. Penetrance estimates are indeed essential for genetic counselling, presymptomatic monitoring or preventive strategies. However, both rarity and co-occurrence with other risk factors make it difficult to estimate the penetrance associated with rare variants. Most common complex disorders indeed result from the combined effect of risk factors each associated with diverse degrees of penetrance.

Alzheimer disease (AD) provides a perfect example of this challenge. AD is a common disorder generally affecting elderly patients. In the vast majority of AD cases, the disease is a complex disorder with a high genetic component (for review, see Nicolas et al.^2^). The ε4 allele of the *APOE* gene (*APOE*-ε4) is the main AD genetic risk factor, both in terms of frequency and of effect size^3^. However, among early onset AD (EOAD) patients, some families exhibit autosomal dominant penetrance due to a pathogenic variant in the *APP, PSEN1* or *PSEN2* genes. In patients negatively screened for these genes, *SORL1* rare protein-truncating variants (PTV) or missense predicted damaging variants were initially identified in 7/29 unrelated EOAD patients with a positive family history of EOAD^4^. In line with *APP, PSEN1* and *PSEN2, SORL1* is involved in the production of A*β* peptides, the aggregation of which being a critical triggering event in AD pathophysiology. However, segregation data of *SORL1* variants remains very scarce, not allowing to assess their mode of inheritance. We showed that such variants are enriched in EOAD cases with a positive family history in a case-control study, with exome-wide significance^5^, and such results were subsequently extended to all AD cases with a clear effect on ages at onset^6–8^. Given the extreme rarity of premature termination codon-introducing variants in controls and very high odds ratios^6^, the question of the penetrance of such variants and a putative use for genetic counseling has been raised. Missense variants with *in silico* predicted deleteriousness have also displayed a strong level of association with the risk of AD, with rarer variants being more strongly associated. However the level of association is of no common measure with what is observed for premature termination introducing variants. Therefore, although some missense variants probably act through a similar loss-of-function effect, evidence suggests that missense variants convey heterogeneous levels of risks. *In vitro* functional analyses are in line with such heterogeneity^9^.

Several approaches have been published to estimate the penetrance of genetic variants. While survival analyses relying on large prospective cohorts with sequencing data available may appear as the gold standard, they find themselves challenged, and thereby impractical, when assessing the penetrance associated with rare variants.

Indeed, they require a large amount of sequencing data and a follow-up period long enough to allow the diagnosis of a sufficient number of cases, which is not yet available. To overcome this limit, methods have been developed to combine data from large prospective cohorts with large case/control studies and evaluate with more accuracy the lifetime risks or penetrance associated with genetic risk variants^3,10^. Another option is to resort to family-based study designs. Indeed, focusing on families where the proband carries a specific variant of interest enriches the dataset in rare variant carriers. However, the downside is the risk of bias resulting from this ascertainment scheme, that could even be compounded by a selection on age at onset. To overcome this issue, methods based on retrospective^11^ or prospective^12,13^ likelihood were developed such as to condition the phenotype observation on the ascertainment process. A more “naive” but effective approach consists in computing the likelihood after exclusion of proband phenotypes^14,15^. All these approaches may be combined with the Elston-Stewart algorithm^16^ in order to deal with the occurrence of missing genotypes, a second major limitation in pedigree datasets. However, these methods remain scarcely used, probably because of their lack of flexibility and their complexity. More recently, Alarcon et al.^17^ proposed a more flexible approach embedded in an expectation-maximization (EM) framework^18^ that alternates between penetrance estimation (M-step) and a belief propagation step to fill in missing genotypes (E-step). It was first used in a context of a monogenic disease, but may be applied to the case of complex diseases and used with parametric or non-parametric survival models combined with proband’s phenotype exclusion. In this paper, we further extend this method to (i) a digenic scenario combining rare and common risk factors and (ii) the integration of previously published data to robustly adjust for the common risk factor. We apply this strategy to assess the penetrance of AD associated with *SORL1* rare loss-of-function (LoF) variants in pedigrees where the AD-affected proband carries such a variant, with a previously published baseline model adjusted for *APOE*-ε4^19^.

## Material and methods

### Participants

#### Recruitment of probands

We first considered all unrelated individuals carrying a *SORL1* rare (allele frequency *<* 1%) non-synonymous variant identified by whole exome sequencing (WES), with a diagnosis of probable AD^20^ among patients referred to the National Reference Center for Young Alzheimer Patients (CNRMAJ, Rouen, France) in the context of a nation-wide recruitment. Ages of onset were ≤ 65 years for previously reported patients^4–6,21^ and *<* 75 for novel patients recruited through the ECASCAD study, whatever their family history and *APOE* genotypes. Diagnoses were based on clinical examination by a neurologist or geriatrician from Memory Clinics and included personal medical and family history assessments, neurological examination, neuropsychological assessment, and neuroimaging. In addition, cerebrospinal fluid biomarkers indicative of AD (as described by Nicolas et al.^22^) were available for 80% of the cases who were selected for WES. All variants were detected in the context of WES performed in a research or in a diagnostic setting in the National Center for Research in Human Genomics sequencing center (CNRGH, Evry, France) using Agilent Sureselect all exons human kits and Illumina sequencing technology, as described before^22^. All patients or legal guardians provided informed written consent for genetic analyses. Patients were recruited following institutional review boards agreements CPP Paris - Ile de France II (RBM-0259) and CPP Ouest 3 (ECASCAD study).

#### Genetic data processing and *SORL1* variants selection

Sequencing data were processed as previously described in Nicolas et al.^22^. After quality control, we excluded patients from non Caucasian ancestry as previously performed in our association studies^5^ as well as those carrying a likely pathogenic or a pathogenic variant in a Mendelian dementia gene including *APP, PSEN1* and *PSEN2*^22^. Among the 1295 remaining exomes of unrelated probands, we extracted those including a germline (allelic ratio [0.25-1], all were eventually heterozygous) rare (allele frequency < 1% in gnomAD non-Finish Europeans^23^), non synonymous and splice region *SORL1* variants (NM003105.6). We considered, as PTV, all nonsense, frameshift and canonical splice site variants, in addition to splice region variants with a demonstrated effect on splicing (one described in Le Guennec et al.^24^ and one unpublished patient blood mRNA analyses). We also considered missense variants that were predicted damaging by all three software tools among Polyphen-2^25^, Mutation Taster^26^, and SIFT^27^, thereafter referred to as Mis3 variants, as candidates for analysis. To lessen the heterogeneity of the set of variants under analysis, we restricted our analysis to the selection of *SORL1* loss-of-function variants, namely (i) high confidence PTVs (i.e. not affecting the last coding exon or 50 bp of the penultimate exon) and (ii) candidate *SORL1* Mis3 variants that demonstrated an *in vitro* loss-of-function effect (Mis3-LoF). More precisely, Mis3-LoF were defined as rare Mis3 variants with at least two *in vitro* assays supporting such a loss-of-function effect and including at least one A*β* peptide measurement. We retained the following three Mis3 as non-ambiguously Mis3-LoF in this conservative approach: c.994C*>*T,p.(R332W); c.1960C*>*T,p.(R654W) (ref nous MedRXiv); and c.1531G*>*C,p.(G511R)^28^. Hereafter and for penetrance estimations, we use the term “LoF” when considering PTV plus Mis3-LoF as a group.

#### Family history and genetic investigations in relatives

Overall, we included 27 probands carrying either a *SORL1* PTV or Mis3-LoF in this study and we contacted all 27 families, in order to extend pedigree information. Clinical status and age at onset for affected relatives or current age for unaffected relatives were obtained at least for siblings and parents of probands and, when possible for aunts, uncles and grandparents. Both parental sides were systematically investigated even when the presence of an affected parent suggested a unilateral transmission. When clinical examination was not possible, a phone interview was performed. Disease status was set as possible or probable AD according to the NINCDS-ADRDA, missing, or unaffected based on available clinical information from medical charts for affected relatives and from available clinical information or phone interviews for unaffected relatives. In addition, we collected blood samples from affected and unaffected relatives with informed written consent, and performed Sanger sequencing to search for the *SORL1* variant segregating in the family and to determine the *APOE* genotype. At the end, we obtained clinical information for 307 relatives with age ≥ 40 years and genotyping information for 45 relatives including 20 carriers of a *SORL1* LoF variant.

### Statistical modeling

To evaluate the age-related penetrance of AD associated with *SORL1* LoF variants, we built a model based on pedigrees. Given the role of common *APOE* alleles in Alzheimer disease, it was essential to account for *APOE* genotypes in this model. However, since the effect of *APOE* alleles has already been deeply studied through large prospective cohort studies, we relied on published results to define the *APOE*-adjusted baseline of our survival model^19^. Given the rarity of *SORL1* variants, we make the assumption that this population-based baseline models Alzheimer disease risk for non carriers of *SORL1* LoF variants. Then our model combines this literature-based baseline for non carriers of *SORL1* LoF variants with an age dependent proportional effect for *SORL1* LoF variant carriers computed from our pedigrees (Figure 1). Since the pedigrees include relatives with missing genotypes, we applied an Expectation-Maximisation (EM) algorithm to take full advantage of the available information on the whole pedigree. This algorithm alternates until convergence between replacing unknown genotypes by individual weights (genotype posterior distributions for each individual) based on current age-related penetrances (E-step) and updating the age-related penetrances for each possible genotype based on observations and previously computed individual weights (M-step).

**Figure 1:**
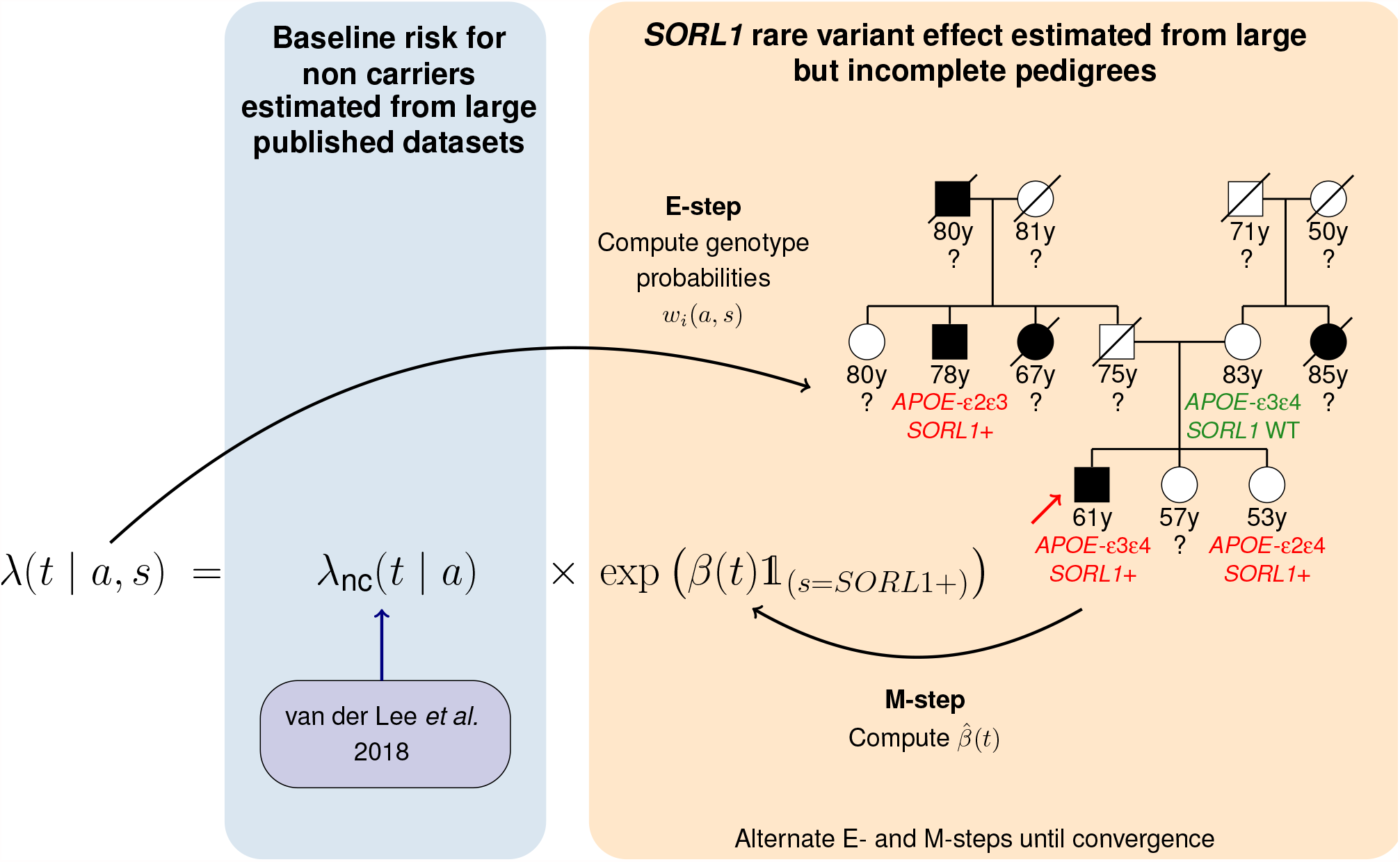
Summary of the method. *λ*(*t* | *a, s*) refers to the instantaneous risk to develop the disease depending on age *t* and genotype (*a, s*) ∈ *APOE* × *SORL*1. *λ*_nc_(*t* | *a*) refers to the specific instantaneous risk associated with non carriers of *SORL1* variant of interest stratified on *APOE* genotype and derived from the Rotterdam study^19^. *β*(*t*) refers to the additional effect of the *SORL1* variant. E/M-steps refer to Expectation/Maximization steps. *w*_*i*_(*a, s*): individual weight updating at each E-step iteration referring to posterior probability distribution of individual *i* for combined genotype (*a, s*); y: years; *SORL1* +: carrier of the variant of interest in *SORL1* gene; *SORL1* WT: wild type for *SORL1* (non carriers of the variant of interest); ?: unknown genotype. The red arrow indicates the proband.

#### Notations

Let *n* be the total number of subjects. Let *Y*_*i*_ be the age at disease onset for subject *i* and *C*_*i*_ the censoring time such that *T*_*i*_ = min(*Y*_*i*_, *C*_*i*_) is the observed time associated with the status *δ*_*i*_ equaling 1 if *T*_*i*_ = *Y*_*i*_ and 0 otherwise. In other words, *δ*_*i*_ indicates if the subject *i* has developed AD at age *T*_*i*_.

We want to estimate the penetrance function *F* (*t* | *a, s*), which gives the probability that an individual develops the disease before age *t* given their *APOE* status *a* and *SORL1* status *s*:

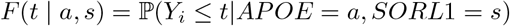

As usually done in survival analysis, the penetrance function is modeled through the survival function *S*(*t* | *a, s*) which models the time to disease onset:

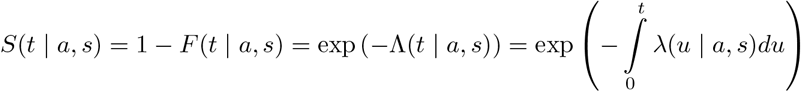

where *λ*(*t* | *a, s*) denotes the hazard function corresponding to the instantaneous risk of developing AD at time *t* conditional on not having developed it before for individuals of *APOE* status *a* and *SORL1* status *s*; and similarly, Λ(*t* | *a, s*) denotes the cumulative risk until time *t* for these individuals. This hazard function takes the following parametric form:

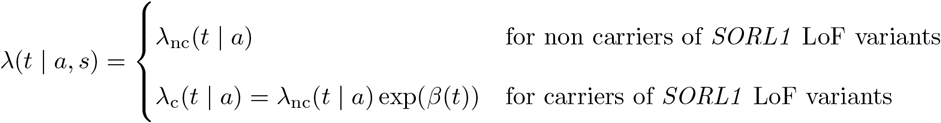

#### Estimating *λ*_nc_(*t, a*)

In this equation, the non carrier hazard rate *λ*_nc_(*t* | *a*) is a piecewise constant function designed to match exactly the *APOE*-specific penetrance estimates derived from age 65 to age 100 in the general population by van der Lee et al.^19^ (see supplementary Table 3 of van der Lee et al.^19^ and supplemental information for detailed computation). As in the original publication, cut offs are located every 5 years from 65 years-old up to 95 years-old. To account for the accumulation of risk before 65 years-old, the first age category spans from 40 to 65 years-old. To compensate for the rarity of *APOE*-ε2 allele, *APOE* genotypes are modeled through the number of *APOE*-ε4 alleles and therefore averaged into three categories: ε4ε4 carriers, ε4 heterozygous carriers and ε4 non-carriers.

#### Estimating *β*(*t*)

Regarding the hazard function for carriers, we also assume *β*(*t*) is piecewise constant over time and most importantly that this additional age-dependent effect of *SORL1* LoF variants does not depend on *APOE*. Because genotype information is not available for most of the subjects in our cohort, we estimated *β*(*t*) using an EM algorithm for the estimation procedure^18^. This iterative algorithm, alternating E- and M-steps to maximize the likelihood of the model, is efficient when the information is partially missing (here the *APOE* and *SORL1* genotypes). At the E-step, we computed individual weights *w*_*i*_(*a, s*) corresponding to the posterior probability that subject *i* should carry genotype {*a, s*} according to current model estimates. At the M-step, we estimate the *β*(*t*) coefficients using weights obtained at the E-step. We denote respectively by 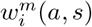 and 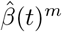 the values of *w*_*i*_(*a, s*) and 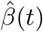 at the *m*^th^ iteration of the EM algorithm.

#### E-step

Denote by 𝒢 the ensemble of the 36 possible *APOE* = *a* × *SORL*1 = *s* genotype combinations. This set accounts for every *APOE* alleles and differentiate paternal from maternal alleles for the purpose of genotype propagation through pedigrees.

The E-step consists in updating for each subject *i* and each genotype combination {*a, s*} ∈ 𝒢 the posterior probability *w*_*i*_(*a, s*) that subject *i* should carry genotype combination {*a, s*} conditional on **ev** the evidence defined as the probability of observing the data according to the genotype and 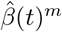, the last estimation of *β*(*t*).

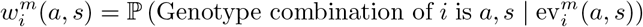

These probabilities are computed using an *ad hoc* C++ implementation of belief propagation in pedigrees (`bped`) extended from Alarcon et al.^17^ to the specific case of *APOE* × *SORL*1 genotypes. Arguments provided to `bped` are the pedigree structure and the *n* × #𝒢 matrix of evidence with:

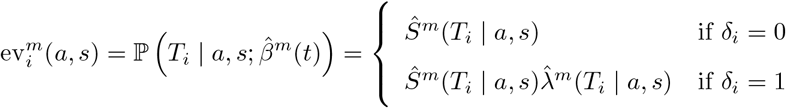

Of note, the evidence and thus posterior probabilities are set to 0 for all genotype combinations {*a, s*} discordant with available information.

#### M-step

We assume that *β*(*t*) is piecewise constant over *J* intervals with ∀*j* ∈ {1, …, *J* }, *β*(*t*) = *β*_*j*_ if *t* ∈ [*τ*_*j*−1_, *τ*_*j*_[, *τ*_0_ = 0 and *τ*_*J*_ = +∞. The M-step consists in estimating *β*_*j,j*∈{1,…,*J*}_ that maximizes the likelihood of our data. Because of unobservable genotypes, we instead maximized 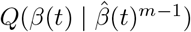 the expected value of the log-likelihood function with respect to the current conditional distribution of unobserved genotype combinations {*a, s*} given evidence and the current estimates of the parameters 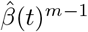:

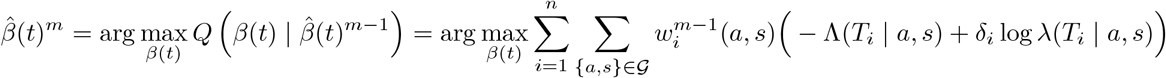

Besides, note that because the additive *SORL1* effect only appears in the survival and hazard function of *SORL1* carriers, the sum on 𝒢 simplifies into a sum on 𝒢_1_, the subset of genotype combinations that include a *SORL1* LoF variant. In the end the parameters 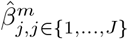 that maximize the *Q* function are straightforwardly obtained by solving for each 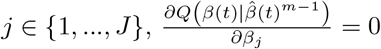 such that:

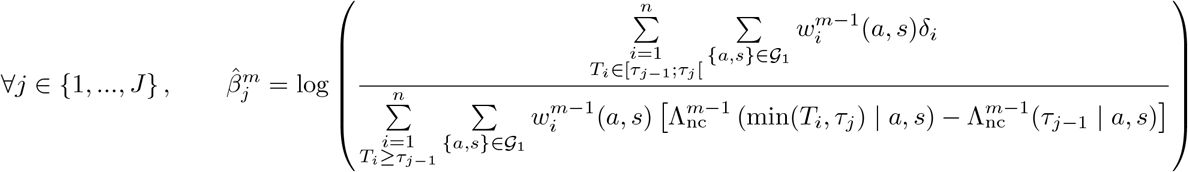

In other words, the hazard ratio exp 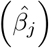 associated with *SORL1* LoF variants on the *j*^*th*^ interval may be interpreted as the ratio of the subjects developing the disease within the time-interval *j* over the time-subjects at risk during this time interval taking into account the *APOE* effect.

#### Correction for ascertainment bias

To tackle the important issue of ascertainment bias resulting from the recruitment of early onset carriers of *SORL1* LoF variants as probands, we took their genotype information into account at each E-step but discarded proband phenotype information (observed time, disease status) at each M-step as in Alarcon et al.^17^. Moreover, since only few individuals’ observation time were greater that 85 years of age, we censored data at 85 years-old.

#### Model choice

To validate the choice of {*τ*_0_, …, *τ*_*J*−1_} for the piecewise modeling of the effect of *SORL1* LoF variants *β*(*t*), several cut-offs were envisaged and the different models were compared based on a Bayesian Information Criterion (BIC) measure defined as:

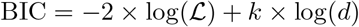

where log(ℒ) is the log-likelihood of the model returned by `bped` computed using the sum-product algorithm from the evidence probability over all possible genotypes, *k* is the number of parameters and *d* the number of subjects who experienced the event of disease onset. The model with the best fit is the one with the lowest BIC.

#### Computation of confidence intervals

Confidence intervals (CI) were computed using a bootstrap strategy of 500 iterations. At each iteration, families were randomly selected with replacement and model parameters were re-estimated on these observations (same family number) to obtain the associated age-related penetrance curves. Then, for each age, we took the 2.5^*th*^ and 97.5^*th*^ quantiles as the boundaries of the 95%CI.

As explained more thoroughly in the supplemental information, uncertainty surrounding *APOE*-adjusted baseline survival curves as reflected by their confidence intervals was taken into account in the bootstrap procedure.

#### Simulation study

We performed a simulation study in order to assess the robustness of our methodology. We proposed several *scenarii* mimicking all the potential biases of our pedigree dataset: age of probands, missing genotype patterns, unbalanced maternal/paternal ratio information in pedigrees as well as unobserved family history effect. We varied simulation parameters and estimated their impact on bias and variance associated with estimates. For each *scenario*, we compared the methodology with and without discarding proband phenotype information. A detailed description of the simulation study is available in supplemental information.

## Results

In order to assess penetrance of rare *SORL1* LoF variants in pedigrees with missing genotypes while adjusting for the *APOE* status, we built a piecewise constant hazard model integrating a literature-based baseline for *APOE*-ε4 effect. The additive effect of *SORL1* variants was estimated from pedigrees through an EM algorithm alternating replacement of unknown genotypes by individual weights (genotype posterior distributions for each individual) based on current age-related penetrance (E-step) and update of the age-related penetrance for each possible genotype based on observations and previously computed individual genotype posterior probabilities (M-step) (Figure 1).

### Simulation study

We first aimed at challenging the performances of our algorithm through simulations. We generated artificial datasets under several *SORL1* variant effects as well as several *scenarii* mimicking the potential biases of a pedigree dataset. We varied simulation parameters and investigated their impact on estimation bias and variance (more details provided in Supplemental Information). Our simulation study confirmed that our methodology was robust to various patterns of genotype and phenotype missingness, in particular even in the presence of unbalanced pedigree ascertainment (Figure 2 and Figures S3-S9). Exclusion of the proband phenotype at the M step (hazard ratio computation) reduced efficiently the bias resulting from age-at-onset-based ascertainment. The Bayesian Information Criterion (BIC) was able to discriminate models and identify the best time cut-offs for the piecewise constant function *β*(*t*) (Figure S10). Based on our simulations study, we concluded that the model might lead to an overestimation of the average variant effect when variant effects are not homogeneous across families.

**Figure 2:**
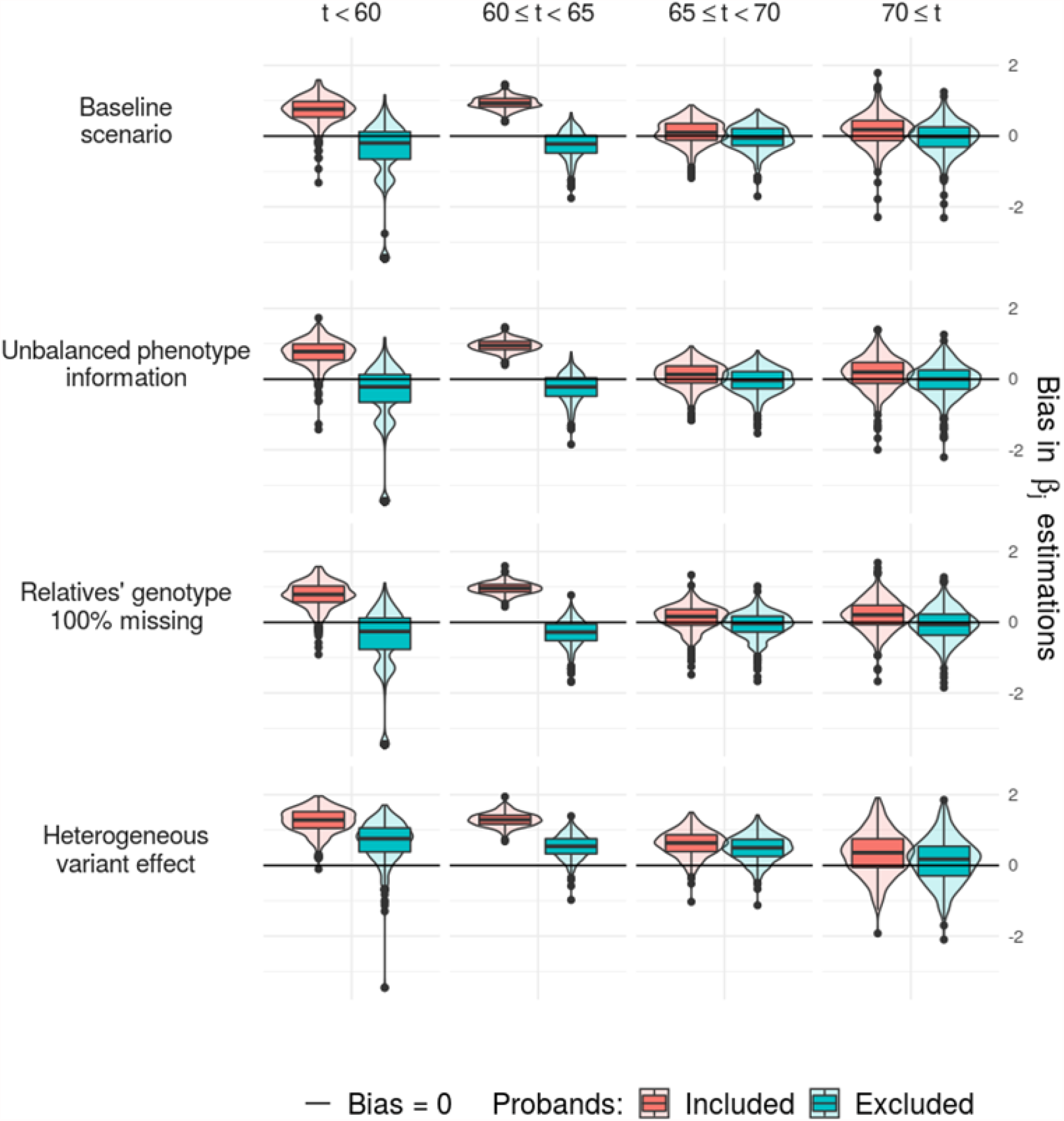
Estimation of bias in our simulation study. Bias was estimated for each of the four constant parameters (in columns) of the piecewise constant *β*(*t*) referring to the additional effect of the *SORL1* variant of interest through 4 *scenarii* of simulation (in rows). Results are provided for probands included and excluded from the analysis during the Maximization-step of the EM algorithm. For the baseline *scenario*, we generated 27 families mimicking what we observed in our dataset in terms of *SORL1* LoF variant effect, ascertainment and available genotypes. Then the model was challenged through three additional *scenarii*: (i) Unbalanced phenotype information: if one of the parents was affected, we removed all the phenotypic information of the parental branch with the unaffected parent. (ii) Relatives’ genotypes 100% missing: we removed all information about relatives’ genotypes. (iii) Heterogeneous variant effect: instead of generating age at onset based on a constant variant effect, we generated age at onset based on a normal distribution of the variant effect with a variance equaling to 1.

### Description of the pedigrees

27 families carried at least one non-ambiguous LoF variant (21 families with one PTV, one family with 2 PTVs, and 5 families with a Mis3 variant with *in vitro* LoF evidence, Mis3-LoF). Although most of the LoF variants were each private to one family, the p.W804* (c.2412G*>*A; PTV) and the p.R654W (c.1960C*>*T; Mis3 with *in vitro* LoF evidence) variants were observed in two and three apparently unrelated families, respectively, even though they were both not observed in the gnomAD database. Among the LoF variants, only 3 were observed at least once in gnomAD (cumulate non neuro AF nfe): p.R332W (c.994C*>*T, minor allele frequency = 1.1 × 10^−5^), p.R866* (c.2596C*>*T; minor allele frequency = 1.9 × 10^−5^) and p.R1207* (c.3619C*>*T; minor allele frequency = 2.2 × 10−5). See Table S1 for a detailed list of variants. In LoF families, proband age of onset varies from 48 to 70 years (mean: 58 years). Two probands (7%) were *APOE*-ε4ε4 carriers (age of onset: 58, 60), 18 (67%) were ε4 heterozygous carriers (age of onset from 50 to 66, mean: 58) and 7 (26%) were ε4 non-carriers (age of onset from 48 to 70, mean: 56). In addition, informative phenotype (disease status and age ≥ 40 years) and genotypes (*APOE* and *SORL1*) were obtained for 307 and 45 relatives, respectively (Table 1). They were collected in both paternal and maternal branches, whatever the apparent disease transmissions (Figure S1). LoF families included a median of 11 informative phenotypes (Q1-Q3: 8.5-17; min-max: 4-24) over 3 generations (Q1-Q3: 3-3; min-max: 2-5) and 2 genotypes (Q1-Q3: 1-4; min-max: 1-6) encompassing data of the proband. Relatives’ genotypes were known (genotyped individual or obligate carrier) for 12 affected carriers (age at onset from 55 to 78, median: 68, Q1-Q3: 64.5-74), 12 unaffected carriers (age from 42 to 95, median: 66, Q1-Q3: 52-68.25), 4 affected non carriers (age at onset: 64, 68, 70 and 75) and 21 unaffected non carriers (age from 37 to 86, median: 68, Q1-Q3: 57-70).

**Table 1:**
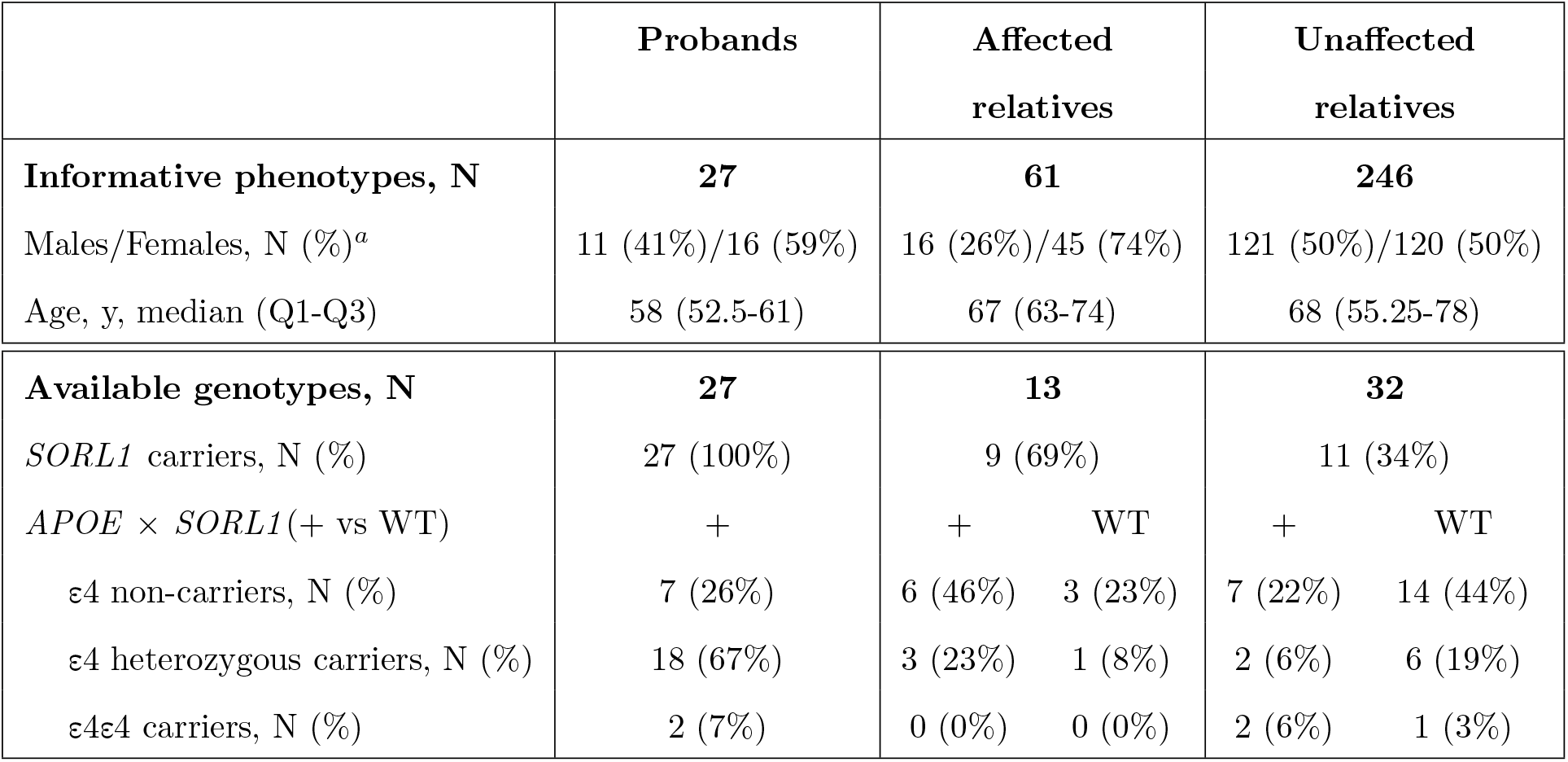
Description of informative phenotypes and genotypes available in the 27 families with a LoF variant. N: number; y: years; Q1: first quartile; Q3: third quartile; WT: wild type. We considered as informative individuals for the phenotype, all those having well-established disease status as well as age at onset or censoring above 40 years. Genotypes were available for 71 (21%) of individuals with informative phenotype as well as for one unaffected relative (censoring age: 37 years, genotype: *APOE*-ε4 heterozygous, *SORL1* WT). a) percentages are given over known gender.

### Genotype probabilities and penetrance estimates

During the EM procedure, genotyping probabilities were attributed to each participant. Final weights added to already known genotype information led to an expected total number of relatives aged ≥ 40 years carrying a *SORL1* LoF variant of 109.6 (36% of relatives aged ≥ 40 years; Figure 3). Table S3 provides detailed estimated genotypes. Based on BIC (Table S2), we retained a piecewise constant *β*(*t*) with cut-offs at respectively 60, 65 and 70 years old (*β*(*t*) = 3.5, 95%CI[1.5; 5.0] for *t <* 60*y*; *β*(*t*) = 6.7, 95%CI[5.1; 8.0] for 60 ≤ *t <* 65; *β*(*t*) = 4.7, 95%CI[3.9; 5.4] for 65 ≤ *t <* 70 and *β*(*t*) = 3.4, 95%CI[2.1; 4.7] for 70 ≤ *t*) leading to an increased age-dependent penetrance for carriers versus non carriers of *SORL1* LoF variants. Following penetrance estimates after adjustment on *APOE*-ε4 status, we observed a full penetrance at the age of 70 years for ε4/4 carriers only, whereas ε4 heterozygous carriers and ε4 non carriers reached respectively 56% (95%CI [40%; 72%]) and 37% (95%CI [26%; 51%]) penetrance (Figure 4). Of note, full penetrance was reached ten years later for ε4 heterozygous carriers (95%CI = [0.91; 1] by age 80) and after 85 years old for ε4 non carriers (95%CI = [0.81; 1] by age 85).

**Figure 3:**
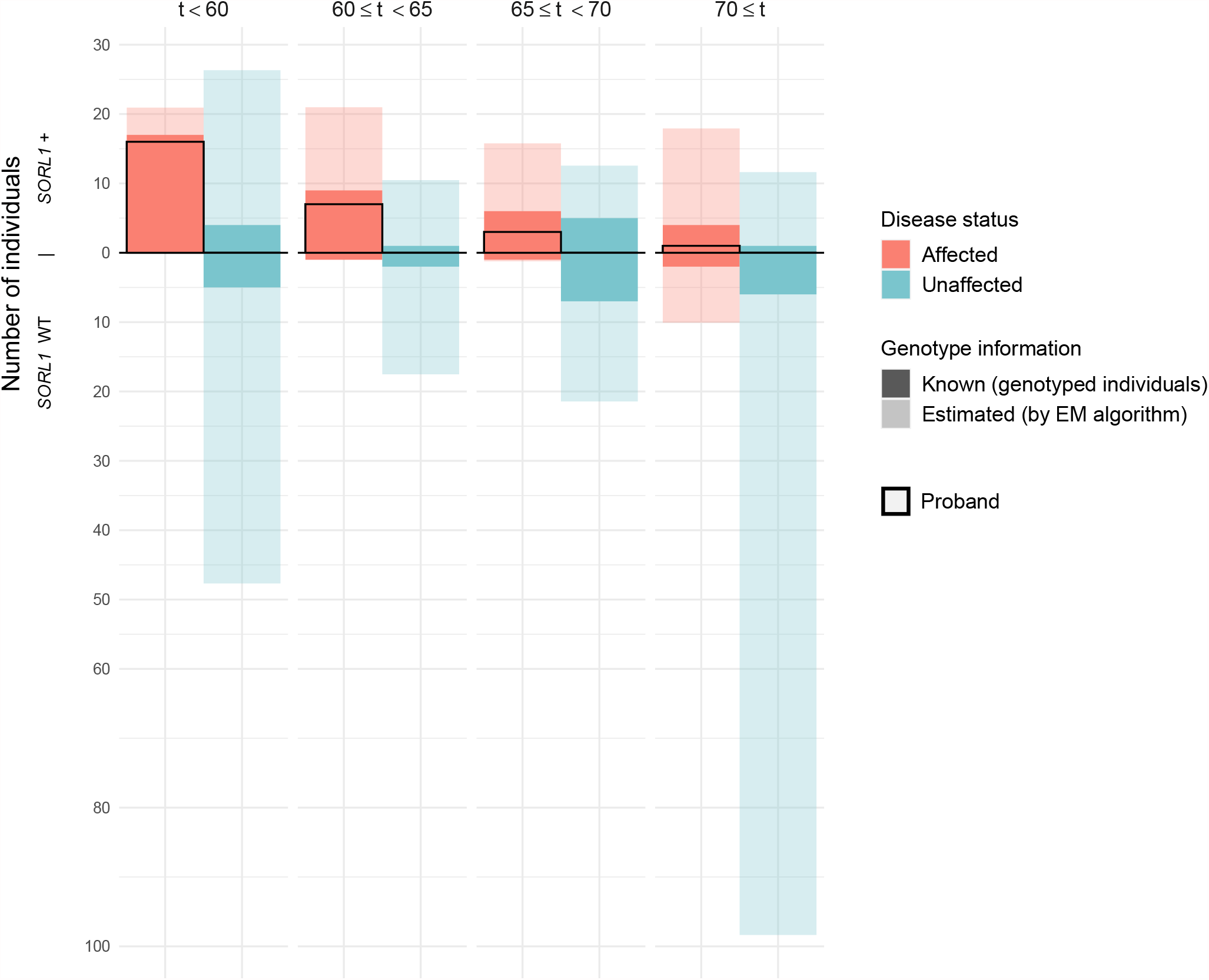
Number of affected and unaffected individuals according to their age and their *SORL1* status after genotyping imputation. This graph was obtained for all individuals with age of onset or censoring above 40 years. It represents the number of carriers (upper part) and non carriers (lower part) of *SORL1* LoF variant according to their disease status and age intervals (age of onset for probands and affected relatives and censoring for unaffected relatives). Transparency differentiates available genotypes (already known, including those of probands) from those estimated at the end of the algorithm.

**Figure 4:**
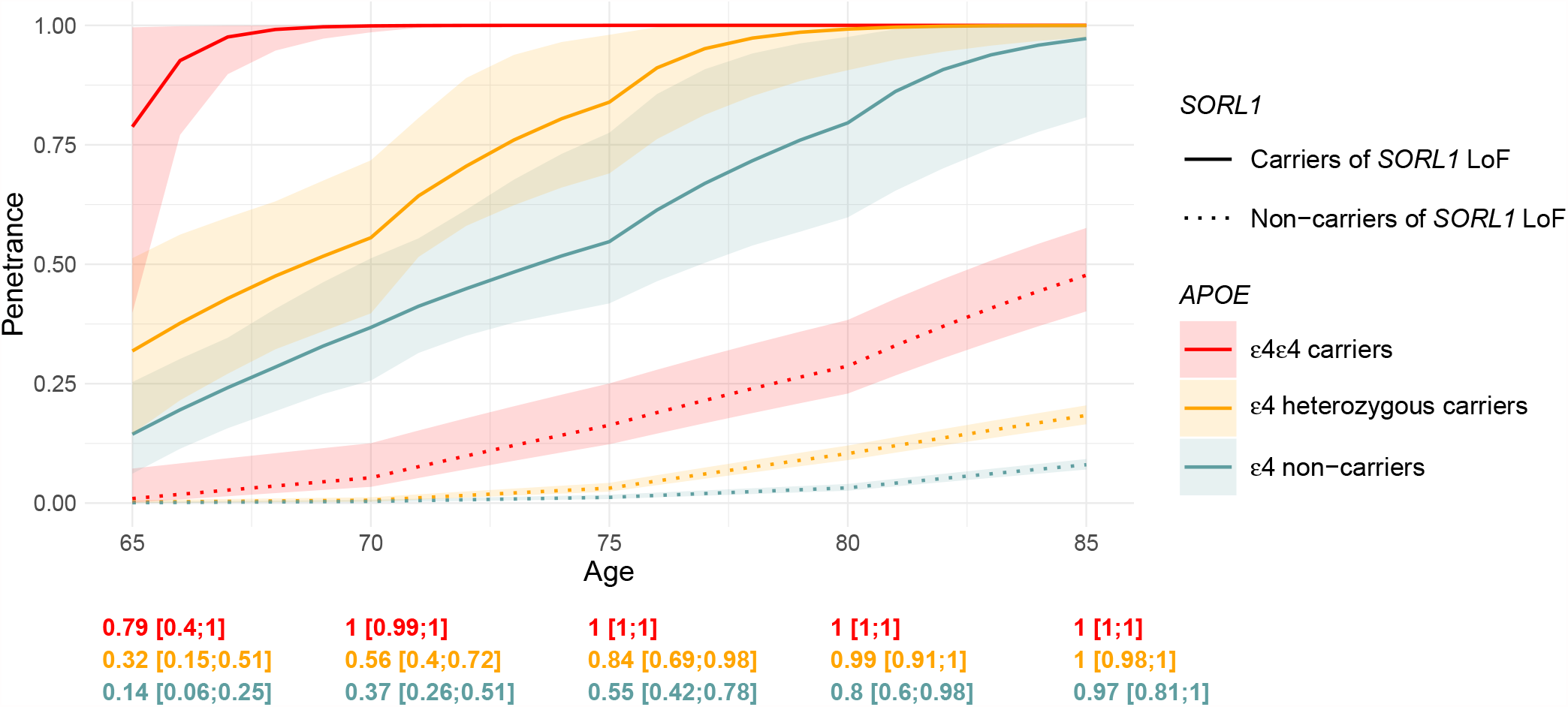
Age dependent penetrance for carriers and non carriers of a *SORL1* LoF variant. The penetrance is displayed with its 95% confidence interval according to the number of *APOE*-ε4 allele from 65 to 85 years of age. Curves for non carriers and their confidence intervals were obtained from our estimation of *λ*_nc_(*t* | *a*) based on the Rotterdam study^19^. Data from pedigrees were censored at 85 years. Confidence intervals for *SORL1* LoF variant carriers were obtained from 2.5th and 97.5th quantiles of 500 bootstrap iterations. Penetrance values at 65, 70, 75, 80 and 85 years of age, for carriers of *SORL1* LoF variant are displayed below the figure.

## Discussion

We investigated the age-related penetrance of AD associated with *SORL1* LoF variants, using a family-based approach and adjusting for the common risk factor *APOE*-ε4. We proposed to model the risk conferred by both factors through a piecewise constant hazard model. As expected, our model indicates that *SORL1* LoF variants confer a high risk before 65 years, and thus are strongly associated with EOAD^6,8^. The risk remained high until 70 years, which is compatible with an association of such a variant with late onset AD (LOAD) among younger patients. However, our results suggested that the penetrance associated with *SORL1* LoF variants should be interpreted in light of the carrier’s *APOE* genotype. Indeed, in our model, the penetrance for *SORL1* LoF carriers was complete by age 70 only among ε4ε4 carriers, whereas the curve for *SORL1* LoF and ε4 heterozygous carriers reached complete penetrance ten years later and even later for ε4 non-carriers.

*APOE*-ε4 has already been investigated as a modifier of AD age at onset regarding the specific *PSEN1* p.E280A pathogenic variant and in APP V717I families^29,30^. However, these variants retained full penetrance before the age of 65 regardless to the *APOE* genotype. Because most pathogenic variants in the *APP, PSEN1* or *PSEN2* genes confer full penetrance before 65 years by themselves, if not earlier, the *APOE*-ε4 effect is difficult to observe for most of these variants and, thus, *APOE* genotypes are not a clinically meaningful information upon genetic counseling in such families. In contrast, *SORL1* LoF variants are less penetrant and thus *APOE*- ε4 substantially modifies ages at onset for carriers of such a variant, suggesting that presymptomatic testing of *SORL1* LoF variants should not be proposed regardless of *APOE* genotyping even though such digenic presymptomatic testing is not usual in clinical genetics consultations and may be difficult to understand. Here, we used stringent criteria for restricting our analysis to a set of variants with homogeneous effect (PTV + Mis3-LoF), in accordance with our simulation study. Indeed, since Mis3 variants were defined solely through bioinformatics prediction tools, we can expect that there is some diversity of the variants’ effects on SorLA function towards A*β* secretion^9,31^. Some of the predicted deleterious variants may remain neutral and some other variants might be associated with an intermediate level of loss of SorLA function regarding A*β* secretion. In our study, we generated familial data based on our nation-wide recruitment of families that is probably among the largest datasets on *SORL1* LoF families. Despite this huge effort based on the sequencing of a large number of probands with a high level of evidence for an AD diagnosis, first, and individual family extension, second, the number of families remained limited, especially after restriction to the families with a non-ambiguous LoF variant. However, the EM approach allowed us to reach sufficient power to provide meaningful estimates. In this study, we relied on the EM framework for penetrance calculation in incomplete pedigrees, first used by Alarcon et al.^17^ and extended to our digenic problematic (E-step extension) and parametric modeling (M-step changes). We finally incorporated it into a 2-step methodology also including a baseline estimation from published data. The major interest for this methodology is the combination of several modules that all may be independently modified and improved for further family-based analyses and should open the door to the analyses of (often incomplete) pedigrees, whatever the disease. Moreover, our model might be further developed to better take into account the following points and thus help refining estimations and apply to other diseases: controlling for ascertainment biases in other ways than removing probands phenotypes information, applying other strategies for cut-off piecewise constant functions, adjusting for family clusters or other genetic factors and taking death as a competing risk.

### The control for ascertainment bias

As probands were selected based on age at onset, we proposed to exclude their phenotypes from model parameters computation (M-step) in order to correct for resulting putative ascertainment bias. Our simulation study showed that this method standardly used in family analyses efficiently helps to correct for this bias, but might lead to an underestimation of parameters at the beginning of the curve. Another approach, based on raking estimators, and more often used in survey analyses^32^, could be developed to assess if such a strategy could better handle this ascertainment bias by incorporating auxiliary information in the procedure and thus adjust for family recruitment probabilities.

### The choice of optimal cut-offs in piecewise constant function

The effect associated with the *SORL1* LoF variants was modeled using a simple but interpretable piecewise constant function. Because we did not know how the instantaneous risk associated with variants changes over time, we compared several models including different cut-offs and selected the best model based on BIC. A possible extension of this method is the repetition of this procedure through bootstrap iterations and a higher quantity of candidate cut-offs resulting in smoother curves^33^. However, non parametric and semi-parametric models like for example the Cox’s model^34^ may be also good candidate models conditionally to their ease of interpretation.

### Adjustment for family cluster

For the sake of simplicity, we considered that individuals were independent conditionally to their *APOE* × *SORL1* genotypes. However, as shown by our simulation study, inter-family heterogeneity may lead to misestimation of model parameters. Stratified models and then frailty survival models were developed to handle this kind of data^35^. The latter proposes to incorporate random term(s) in the model mimicking an intra-familial correlation and thus reduce the part of variability associated with *SORL1* variants effect.

### The death as competing risk

Here, we have chosen to consider death as censoring, meaning that death and disease onset are independent. Indeed, there is no demonstrated increased risk of death linked to *SORL1* LoF variants. In contrast, *APOE*-ε4 is related to a higher risk of mortality independently of AD^36^. Since this was already taken into account in published data we used for baseline risk according to *APOE*, we did not need to adjust a second time for it^19^. However, in future applications of this methodology to the case of other diseases, the M-step may easily incorporate models developed in a competing risk framework^37^.

In our model, we adjusted on the number of *APOE*-ε4 alleles given the frequency of this allele and its moderate to high impact. In addition to *APOE* and *SORL1*, other types of variants can influence AD risk. Common variants identified in GWAS and rare variants identified in exome sequencing studies, mainly in the *ABCA7* and *TREM2* genes, can thus modulate the individual risk as computed using *SORL1* and *APOE* only. Recently, polygenic risk scores (PRS) have been developed, gathering the individual small effect of multiple GWAS SNPs.

Although there is not a unique method to build PRS, including the selection of variants that can range from a restricted number of SNPs associated with AD after Bonferroni correction to a large number SNPs with an association p-value below 0.5, the effect of PRS as modulators of the *APOE* genotype has been assessed^19^. Adjusting the effect of rare *SORL1* variants on that PRS-adjusted *APOE*-ε4 is thus an interesting perspective, conditional to the feasible extension of the E-step to such a multigenic model, i.e. allowing for the imputation of multiples SNPs for PRS estimation in relatives without drastically increasing computational cost. However, given the large risk load already conferred by *APOE* and *SORL1*, we do not expect that PRS may play a clinically-relevant role in *SORL1* +*APOE*-ε4 carriers. This might however be different in case of co-occurrence of an *ABCA7* or a *TREM2* risk variant. Indeed, odds ratios of rare deleterious variants of either gene are in the order of magnitude of 2.5-4, and even higher for the extremely rare *TREM2* PTVs^21^, and lower for the R62H *TREM2* variant^38^. Thus, an oligogenic model taking into account these genes would be welcome. However, the extreme rarity of carriers of unambiguously deleterious/associated *TREM2, ABCA7* and *SORL1* variants make such an oligogenic model challenging to assess with sufficient power.

In conclusion, we propose here digenic penetrance estimations for *SORL1* LoF variants together with *APOE*-ε4 alleles. We used a family-based approach that demonstrated to be powerful to provide meaningful estimations. Our recruitment was likely closer to that of genetic counseling requests in families than to the general population, and our estimates may thus be taken into account when an asymptomatic relative may request information following the identification of a *SORL1* LoF variant found in a proband. We consider that *SORL1* presymptomatic testing should not be performed regardless of *APOE* genotyping, given the large modifying effect in these families. Our estimations may be also useful for elaborating preventive clinical trials based on similar recruitment criteria. Further work remains necessary to estimate the penetrance of other *SORL1* missense variants as well as rare variants in other risk genes, such as *TREM2* and *ABCA7*. Finally, an integrative model is warranted in exceptional cases of co-occurrence of rare deleterious variants in multiple genes, together with the *APOE* genotype, that may additionally be PRS-adjusted.

## Supplemental Data

Supplemental Data include supplemental methods, detailed results from the simulation study, eleven figures and three tables.

## Acknowledgments

This work was supported by grants from Fondation pour la Recherche Médicale (Equipe FRM DEQ20170336711), Fondation Alzheimer (ECASCAD study), France Alzheimer association (AAPSM2019 – grant #1957) and CNRMAJ. CS is supported by the Fondation pour la Recherche Medicale (ARF201909009263). This work is a collaboration between CEA-DRF-Jacob-CNRGH-CHU de Rouen.

## CNRMAJ collaborators, by city

Ales: Julia Nivelle; Amiens: Daniela Andriuta, Mélanie Barbay, Olivier Godefroy, Alexandre Perron; Angers: Valérie Chauvire, Frédérique Etcharry-Bouyx, Virginie Pichon; Arles: Marie De Verdal; Bayonne: Guillaume Ballan; Beaune: Fabienne Contegal-Callier; BesanÇon: Sophie Haffen, Eloi Magnin, Alice Voilly; Bordeaux: Sophie Auriacombe, Chloé Gregoire, Brice Laurens, Vincent Planche; Brest: Amélie Leblanc; Caen: Pierre Branger, Julien Cogez, Sophie Dautricourt, Olivier Martinaud, Vincent de la Sayette; Clermont-Ferrand: Didier Deffond, Elsa Dionet; Colmar: Pierre Anthony, Geoffroy Hautecloque, François Sellal; Dijon: Yannick Bejot, Benoit Delpont, Mathilde Graber, Julien Gueniat, Catia Khoumri, Sophie Mohr, Christel Thauvin; Grenoble: Olivier Moreaud, Mathilde SauvÉe; Lille: Pascaline Cassagnaud, Yaohua Chen, Charlotte Crinquette, Vincent Deramecourt, Céline Derollez, Thibaud Lebouvier, Marie-Anne Mackowiak, Aurélien Maureille, Florence Pasquier, Adeline Rollin-Sillaire; Limoges: Leslie Cartz-Piver, Philippe Couratier, Benjamin Dauriat; Lyon: Bernard Croisille, Maïté Formaglio, Hélène Mollion, Elisabeth Ollagnon Roman; Marseille: Mathieu Ceccaldi, Lea Corneille, Mira Didic, Boris Dufournet, Stephan Grimaldi, Claude Gueriot, Lejla Koric; Montpellier: Karim Bennys, Giulia Diemert, Audrey Gabelle, Pierre Labauge, Anaïs Lippi; Cecilia Marelli, Cédric Turpinat; Mulhouse: Emmanuelle Ginglinger; Nancy: Thérèse Jonveaux; Nantes: Claire Boutoleau BretonniÈre, Hélène Courtemanche, Nathalie Wagemann; Nice: Annabelle Chaussenot; Nimes: Giovanni Castelnovo, Camille Heitz, Lila Sirven Villaros; OrlÉans: Hélène-Marie LanoiselÉe, Christophe Tomasino; Paris: Stéphanie Bombois, Emmanuel Cognat, Astrid DeliÈge, Julien Dumurgier, Lorraine Hamelin, Claire Hourregue, Julien Lagarde, Isabelle Le Ber, Camille Noiray, Claire Paquet, Pauline Rod-Olivieri, Carole RouÉ-Jagot, Dario Saracino, Marie Sarazin; Perpignan: Anaïs Dutray, Snejana Jurici, Laurène Van Damme; Poitiers: Gwenaël Le Guyader; Quimper: Philippe Diraison; Reims: Martine Doco Fenzy, Céline Poirsier; Rennes: Serge Belliard, Florence Demurger, Anne Gainche-Salmon, Cezara Hanta; Rouen: Aude Doan, Didier Hannequin, Clémence Hardy, Morgane Lacour, Alexandre Morin, Gaël Nicolas, David Wallon, Aline Zarea; Saint Brieuc: Olivier Vercruysse; Saint Etienne: Jean-Claude Getenet; Strasbourg: Frédéric Blanc, Benjamin Cretin, Hélène Durand, Nathalie Philippi; Toulouse: Marie Benaiteau, Jasmine Carlier, Emilie Milongo Rigal, Jérémie Pariente, Marie Rafiq, Camille Tisserand; Tours: Anna-Chloé Balageas, Emilie Beaufils; Villeurbanne: Christine Champion, Pierre Krolak-Salmon .

