## Supplemental Information for "Penetrance estimation of *SORL1* loss-of-function variants using a family-based strategy adjusted on *APOE* genotypes suggest a non-monogenic inheritance"

### 1 Supplemental material and methods

#### 1.1 Estimating $\lambda_{\mathbf{nc}}(t \mid a)$, the piecewise constant hazard model for the baseline age-related penetrance adjusted on *APOE* genotype

Let $\lambda_{\text{nc}}(t \mid a)$ be the piecewise constant hazard function for non-carriers of a *SORL1* LoF variant, depending on age $t$ and *APOE* genotype $a$. Since *SORL1* LoF variants are extremely rare, general population is an acceptable proxy for non-carriers. Thus we estimated parameters for $\lambda_{\text{nc}}(t \mid a)$ such that $F_{\text{nc}}(t \mid a)$ matches with $P(t \mid a)$, the penetrance estimated every 5 years from 65 years-old up to 95 years-old by van der Lee et al.[1] in general population (see supplementary Table 3 of van der Lee et al.[1]).

Published data did not include case with disease onset before 60 years old, suggesting that $\forall t \leq 60,\ P(t \mid a) = 0$. To account for the accumulation of risk before 65 years-old, in carriers of *SORL1* LoF variant, we propose to cut time into the following intervals $\mathcal{I} = \Big\{ ]40;65],\ ]65;70],\ ]70;75],\ ]75;80],\ ]80;85],\ ]85;90],\ ]90;95],\ > 95 \Big\}$ with:

$$\lambda_{\text{nc}}(t \mid a) = \begin{cases} 0 & \text{if } t \in [0;40] \\ \exp(\alpha_{a,k}) \text{ with } \alpha_{a,k} \sim \mathcal{N}(\mu_{a,k}, \sigma^2_{a,k}) & \text{if } t \text{ is in the } k^{th} \text{ interval of } \mathcal{I}\text{: } ]t_{k-1};t_k] \end{cases}$$

such that $F_{\text{nc}}(t \mid a)$ is not null on $]40, 65]$. Instead of estimating $\alpha_{a,k}$ from published mean penetrance estimation, we propose to estimate $\mu_{a,k}$ and $\sigma_{a,k}$ from published confidence intervals in order to integrate uncertainty linked to *APOE* into our estimations.

For each *APOE* genotype $a$ and $\forall k$, $\mu_{a,k}$ and $\sigma_{a,k}$ are estimated by:

$$\mu_{a,k} = \frac{\alpha^{(L)}_{a,k} + \alpha^{(U)}_{a,k}}{2} \qquad \text{and} \qquad \sigma_{a,k} = \frac{\alpha^{(U)}_{a,k} - \alpha^{(L)}_{a,k}}{2 \times 1.96}$$

where $\alpha^{(L)}_{a,k}$ (respectively $\alpha^{(U)}_{a,k}$) was deduced from $P^{(L)}(t,a)$ (respectively $P^{(U)}(t,a)$), the lower (respectively upper) bound of the 95% confidence interval of published penetrance curve as follow:

$$\exp(\alpha^{(L)}_{a,k}) = \frac{-1}{t_k - t_{k-1}} \times \log\left(\frac{1 - P^{(L)}(t_k \mid a)}{1 - P^{(L)}(t_{k-1} \mid a)}\right) \qquad \exp(\alpha^{(U)}_{a,k}) = \frac{-1}{t_k - t_{k-1}} \times \log\left(\frac{1 - P^{(U)}(t_k \mid, a)}{1 - P^{(U)}(t_{k-1} \mid a)}\right)$$

Of notes, estimates displayed in supplementary Table 3 of van der Lee et al.[1] are rounded to 0 for $P^{(L)}(t = 65 \mid a = \varepsilon3\varepsilon3)$ and for $P^{(L)}(t = 65 \mid a = \text{heterozygous } \varepsilon4)$. To overcome this problem, we proposed to approximate $P^{(L)}(t = 65 \mid a)$ by:

$$1 - \exp\left(-(65-40) \times \exp\left(2 \times \log\left(-\frac{\log(1 - P(t = 65 \mid a))}{65 - 40}\right) - \log\left(-\frac{\log(1 - P^{(U)}(t = 65 \mid a))}{65 - 40}\right)\right)\right)$$

for $a = \varepsilon3\varepsilon3$ and $a = \text{heterozygous } \varepsilon4$ respectively.

Finally, to compensate for the rarity of *APOE* ε2, *APOE* genotypes are modeled through the number of *APOE* ε4 alleles and therefore averaged into three categories: no allele ε4, heterozygous ε4 and ε4ε4 individuals. Of notes, published estimates for non-carriers of ε4 allele are split between ε3ε3 on the one hand and individuals carrying genotypes ε2ε2 or ε2ε3 on the other hand. We recomputed $P(t, \text{no allele ε4})$ using the following formula:

$$\forall t \in \mathbb{R}^+,\; P(t \mid \text{no allele ε4}) = \frac{p_{33} \times P(t \mid \text{ε3ε3}) + p_{22+23} \times P(t \mid \text{ε2ε2 or ε2ε3})}{p_{33} + p_{22+23}}$$

where $p_{33}$ and $p_{22+23}$ are respectively the proportion of *APOE*-ε3ε3 and *APOE*-(ε2ε2 + ε2ε3) genotypes in the data set.

#### 1.2 Computing confidence intervals for penetrance curves

**Uncertainty surrounding *APOE* effect.** Confidence intervals for non carriers of *SORL1* variant are recomputed from parameters $\mu_{a,k}$ and $\sigma_{a,k}$ defined in section 1.1 of supplemental information using the following procedure:

- Repeat 500 times the steps 1 to 4:

  1. Generate $u \sim \mathcal{N}(0, 1)$
  2. $\forall k, \forall a$, compute $\alpha_{a,k} = \mu_{a,k} + \sigma_{a,k} \times u$
  3. Define the function $\lambda_{\text{nc}}(t \mid a)$ from $\alpha_{a,k}$ as in section 1.1 of supplemental information
  4. Compute the penetrance $F_{\text{nc}}(t \mid a) = 1 - \exp\left(-\int_0^t \lambda_{\text{nc}}(u \mid a) du\right)$

- We determined the pointwise 95%CI as the $2.5^{th}$ and $97.5^{th}$ quantiles at each age $t$ over the 500 estimations.

**Uncertainty surrounding both *SORL1* and *APOE* effects.** Confidence intervals for carriers of *SORL1* variant are computed combing a bootstrap strategy of 500 iterations and the strategy described above, by using the following procedure:

- Repeat 500 times the steps 1 to 6:

  1. Randomly select families with replacement (same number of families as in the whole dataset)
  2. Generate $u \sim \mathcal{N}(0, 1)$
  3. $\forall k, \forall a$, compute $\alpha_{a,k} = \mu_{a,k} + \sigma_{a,k} \times u$
  4. Define the function $\lambda_{\text{nc}}(t \mid a)$ from $\alpha_{a,k}$ as in section 1.1 of supplemental information
  5. Compute $\beta(t)$ using EM algorithm incorporating the current dataset and the current $\lambda_{\text{nc}}(t \mid a)$ function
  6. Compute the penetrance $F_{\text{c}}(t \mid a) = 1 - \exp\left(-\int_0^t \lambda_{\text{nc}}(u \mid a) \exp\left(\beta(t)\right) du\right)$

- We determined the pointwise 95%CI as the $2.5^{th}$ and $97.5^{th}$ quantiles at each age $t$ over the 500 estimations.

##### 1.3 Simulation study

We propose to challenge our model in a simulation study. For each *scenario*, we generated 500 independent datasets and apply the model described in the main manuscript on each of them. Data were generated using the following steps:

- Initialisation step:

  Let $N = 27$ the number of families to include in the cohort.

  $n \leftarrow 0$.

- While $n < N$, do:
  - Generate a pedigree structure ($\mathcal{P}^\star$)
  - Generate genotypes by randomly attributing founders *APOE* genotype based on allele frequencies computed from the $11,375$ individuals included in van der Lee et al.[1] (allele frequency for ε2: 0.085; ε3: 0.764; ε4: 0.151). The *SORL1* variant has been randomly attributed to one of the founder. Then, genotype of non-founders were iteratively deduced from their parents with an allele transmission probability of 1/2.
  - Generate phenotypes

    Generate age at AD onset $Y$ according to the hazard function $\lambda^\star(t)$

    Generate censoring time $C \sim \mathcal{N}(65, 15^2)$

    Let $T = \min(Y, C)$, the observed time and $\delta = \mathbb{1}_{Y \leq C}$ the disease status
  - Select proband and its family

    If the third generation of the family includes an affected individual with AD onset before 65 years and carrier of a *SORL1* variant, then this individual is defined as the proband, the family is included in the cohort and $n \leftarrow n + 1$.

- Determine available data in terms of phenotype ($\varphi^\star$) and genotype ($\Theta^\star$). Of note, proband's phenotype and genotype are always known.

We detailed below the choice of $\mathcal{P}^\star$, $\lambda^\star(t)$, $\phi^\star$ and $\Theta^\star$ for the baseline *scenario* and how they were adapted for *scenarii* A, B, and C assessing respectively the impact of the proportion of missing genotypes, the proportion of missing phenotypes and the addition of a familial effect on the estimation, as well as for *scenario* D modifying the number of cut points for the piecewise constant function $\beta(t)$ to assess the performance of the BIC in the choice of the model.

**The baseline scenario**

The pedigree structure ($\mathcal{P}^\star$)

To be congruent with number of available data in our cohort of families with LoF variants, we simulated pedigree of 12 individuals over three generations: four grand-parents (founders, generation 1), maternal (respectively paternal) branch including the mother (respectively father) and a uncle/aunt (generation 2) and the proband with three siblings (generation 3). Considering the censoring distribution, this lead to a mean of 11 informative individuals (age $\geq$ 40) by family (see Table 1 and Figure S2).

The model ($\lambda^\star(t)$)

Age at AD onset $T$ was generated using the inverse transformation method, according to the following model:

$$\lambda^\star(t \mid x, z) = \lambda_{\text{nc}}(t \mid x) \exp(\beta(t) \mathbb{1}_{z=1})$$

where $x$ is the number of ε4 allele and $z$ the indicator of the presence of a *SORL1* variant of interest. We supposed three hypotheses for variants effect $\beta(t)$:

- Hypothesis 0: $\beta(t) = 0 \quad \forall t \in \mathbb{R}^{*+}$
- Hypothesis 1: $\beta(t) = \begin{cases} 3.46 & \text{if } 0 < t \leq 60 \\ 6.67 & \text{if } 60 < t \leq 65 \\ 4.68 & \text{if } 65 < t \leq 70 \\ 3.25 & \text{if } 70 < t \end{cases}$
- Hypothesis 2: $\beta(t) = \begin{cases} 1.78 & \text{if } 0 < t \leq 60 \\ 3.34 & \text{if } 60 < t \leq 65 \\ 2.34 & \text{if } 65 < t \leq 70 \\ 1.63 & \text{if } 70 < t \end{cases}$

Under hypothesis 0, we suppose that the *SORL1* variant has no effect on AD whereas under hypothesis 1, we suppose that the effect is similar to the one we computed through the real data. Finally, we evaluateD the model under a more moderate effect of *SORL1* variants (hypothesis 2).

Available data for the analysis ($\phi^\star$ and $\Theta^\star$)

We suppose that the phenotype is known for all individuals in the cohort.

We suppose that the genotype is known for the proband and for 10% and 20% of its relatives of generation 2 and 3 with $Y \geq 70y$ and $Y < 70y$ respectively, which reflect the reality of our cohort. We suppose that genotype is not available for grand-parents (generation 1).

**Scenario A: assessing the impact of missing genotypes on $\beta(t)$ estimation**

Elements $\mathcal{P}^\star$, $\lambda^\star(t)$ and $\phi^\star$ are unchanged compared with the baseline *scenario*. The proportion of available genotypes varies according to the following possibilities:

- 100% available
- 100% missing (except the proband)
- available for the proband and for 10% of its relatives of generations 2 and 3
- available for the proband and for 50% of its relatives of generations 2 and 3
- available for the proband and for 0% and 10% of its relatives of generations 2 and 3 with respectively $Y \geq 70y$ and $Y < 70y$
- 0% and 30% of its relatives of generations 2 and 3 with respectively $Y \geq 70y$ and $Y < 70y$

**Scenario B: assessing the impact of missing phenotypes on $\beta(t)$ estimation**

Elements $\lambda^\star(t)$ and $\Theta^\star$ are unchanged compared with the baseline *scenario*. The proportion of available phenotypes for proband's relatives varies according to the following possibilities:

- 10% of missingness
- 50% of missingness
- 100% of missingness in one of the paternal or maternal branch (randomly selected)
- 50% of missingness in the parental branch with the less affected cases (both branches were kept if ex aequo)/parent without disease
- 100% of missingness in the parental branch with the less affected cases (both branches were kept if ex aequo)/parent without disease

Since small families may be less impacted by unbalanced missingness between maternal and paternal branches, this *scenario* was assessed for a larger pedigree structure adding three uncles/aunts in both parental branches for a total of 18 individuals.

**Scenario C: assessing the impact of an additional family effect**

Elements $\mathcal{P}^\star$, $\phi^\star$ and $\Theta^\star$ are unchanged compared with the baseline *scenario*. Here we propose to modify the model for generating $T$ according to a new definitions of $\lambda^\star(t \mid x, z)$ including an additional family effect :

$$\lambda^\star(t \mid x, z) = \lambda_{\text{nc}}(t \mid x) \exp(\beta(t)\mathbb{1}_{z=1}) \times \nu$$

where $\nu$ is defined according to the following possibilities:

- $\nu = \exp(u\mathbb{1}_{z=1})$ with $u \sim \mathcal{N}(0, \sigma^2)$, $\sigma = 0.3$
- $\nu = \exp(u\mathbb{1}_{z=1})$ with $u \sim \mathcal{N}(0, \sigma^2)$, $\sigma = 0.5$
- $\nu = \exp(u\mathbb{1}_{z=1})$ with $u \sim \mathcal{N}(0, \sigma^2)$, $\sigma = 1$

} additional effect only affects carriers of the *SORL1* variant of interest

- $\nu \sim \Gamma(1, \theta)$, $\theta = 10$
- $\nu \sim \Gamma(1, \theta)$, $\theta = 1$

} additional effect affects similarly the whole family

Of notes, the baseline *scenario* corresponds to the cases $\sigma = 0$ or $\frac{1}{\theta} \to 0$.

For all *scenarii* above, data were generated under hypotheses 0, 1 and 2 and then analysed using the EM algorithm presented in the main article, for 3 cut-offs (60, 65, 70), when excluding and when including probands' phenotypes.

**Scenario D: assessing the pertinence of BIC for the choice of the cut-points**

Elements for scenario D are unchanged compared to the baseline scenario. Besides hypothesis 1, data were generated under two additional hypotheses including a lower number of cut-points:

- Hypothesis 3: $\beta(t) = 4.08 \quad \forall t \in \mathbb{R}^{*+}$
- Hypothesis 4: $\beta(t) = \begin{cases} 4.74 & \text{if } 0 < t \leq 70 \\ 3.22 & \text{if } 70 < t \end{cases}$

Then, data were analysed using the EM algorithm presented in this article when excluding probands. Several cut-points were tested and BIC was computed for each of them.

### 2 Supplemental Results

#### 2.1 Results from the simulation study

*Scenario A:*

In all *scenarii*, the inclusion of the proband in the analyses lead to a positive bias in $\beta(t)$ estimation before 65 years old. In constrast, the exclusion of the proband corrected for this bias. Results suggested that proportion of available genotype did not impact the bias, but a higher proportion of already known genotype help to reduce the uncertainty of our estimation (see Figures S3 and S4).

*Scenario B:*

In all *scenarii*, missing phenotype does not impact the results, even in unbalanced family design (see Figures S5, S6 and S7).

*Scenario C:*

When adding a random term associated with an heterogeneous variant effect across families or a family effect, the model may lead to an overestimation of $\beta(t)$ even if proband was excluded from the analysis. Besides the non consideration of a random term in the model, the ascertainment based on the age of the proband lead to the inclusion of families whose associated random terms are biased towards higher effect (see Figures S8 and S9).

*Scenario D:*

Whatever the number of cut-offs we considered, over the 500 simulated datasets, the median of BIC was always lower for the model including the right cut-offs (Figure S10). In the datasets generated from model without cut-off ($\beta(t)$ contant over time), the BIC increased with number of cut-offs but the low pairwise difference in BIC suggests that an higher number of cut-offs is not deleterious for the estimation. In contrast, as expected,in the datasets generated from model including cut-offs, simulation results suggested that do not include the original cut-off(s) in the estimation lead to a higher BIC. Thus, BIC is an acceptable criterion for the choice of our model.

*Genotyping probabilities*

From the baseline scenario (under hypothesis 1), we showed that the sum of posterior genotyping probabilities may correctly estimate the proportion of carriers of the variant of interest (Figure S1).

#### 3 Supplemental Figures

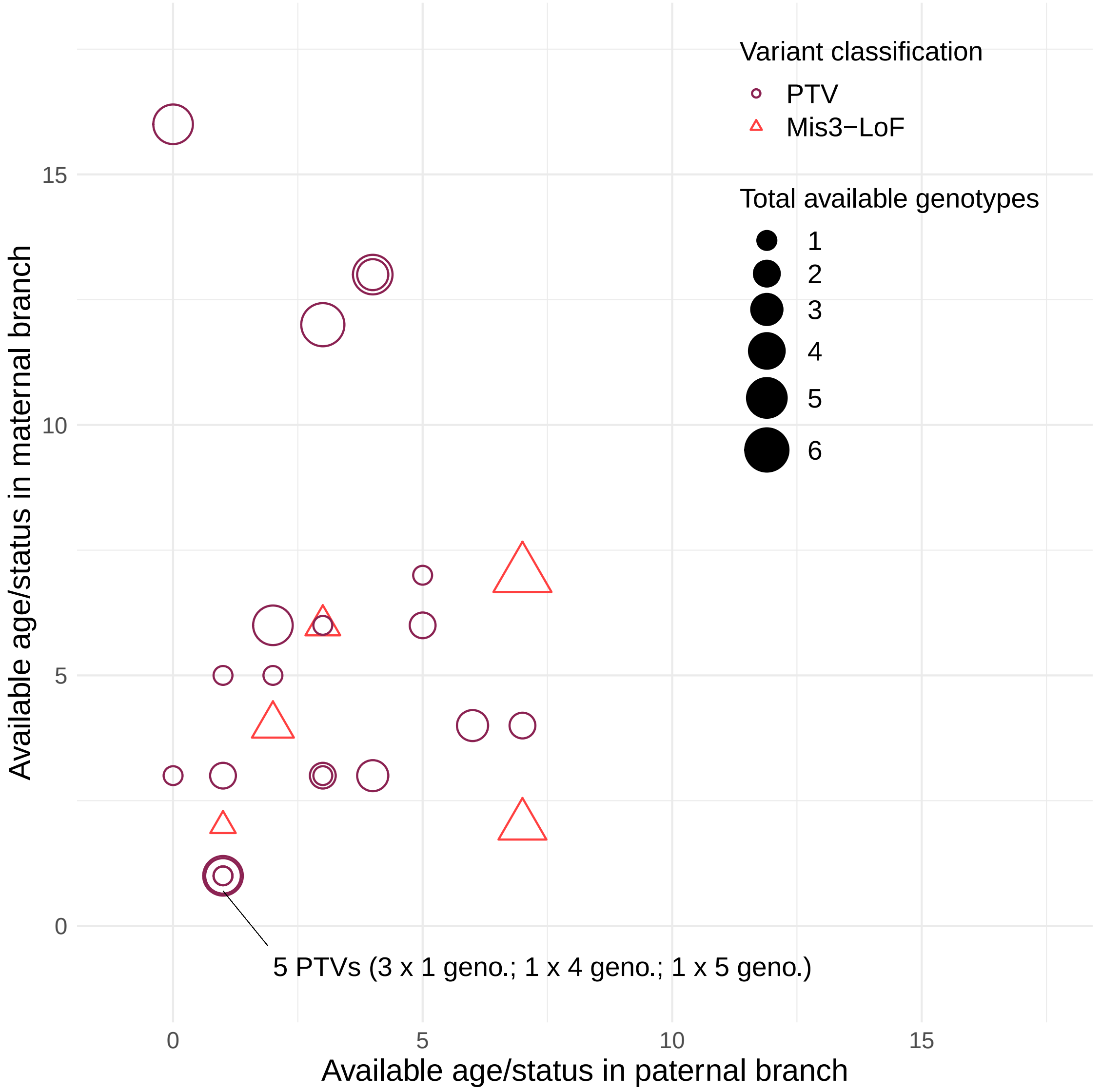


Figure S1: Number of informative phenotypes available in maternal versus paternal branch over the 27 LoF families.
Each point represents a family. Point shape/color corresponds to the variant classification and point size corresponds to the total number of genotypes (*APOE* and *SORL1*) available in the family, including the proband genotype. The X-axis (respectively the Y-axis) represents the number of individuals in the paternal (respectively maternal) branch with informative phenotype (i.e. status and age $\geq$ 40 years old). Of notes, X- and Y- axes do not count probands, siblings, nieces/nephews nor those that are not related by blood.

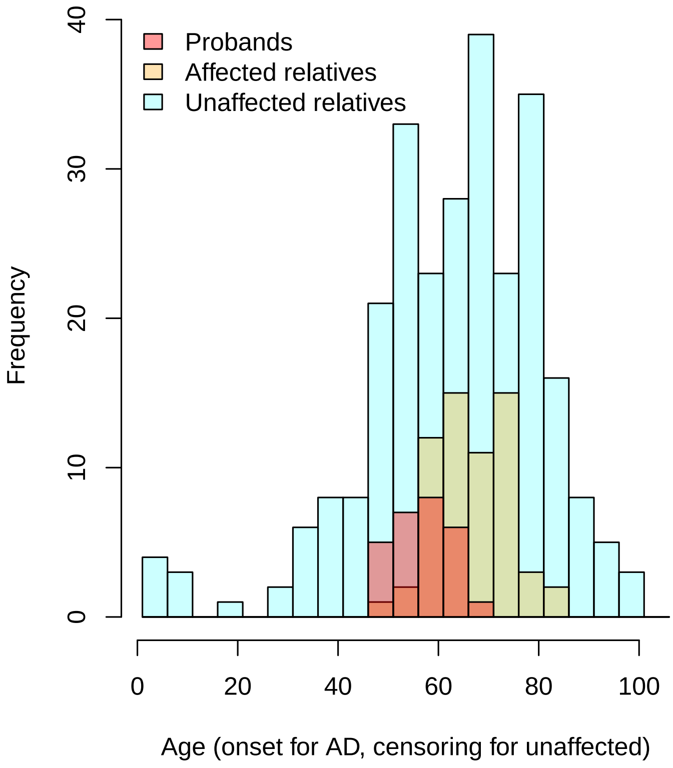


Figure S2: Age distribution in LoF families.

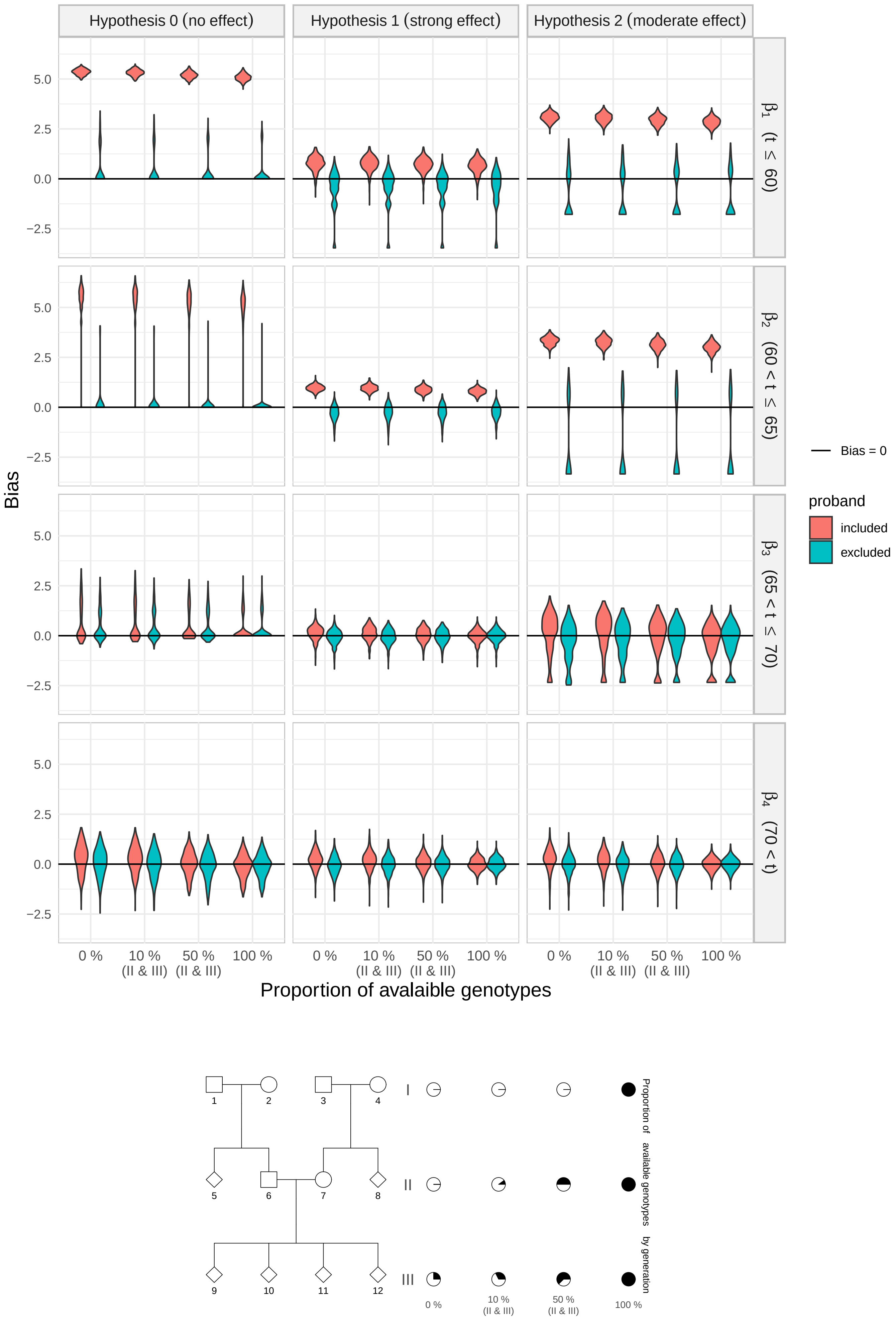


Figure S3: Results of simulations in scenario A when missingness does not depend on age

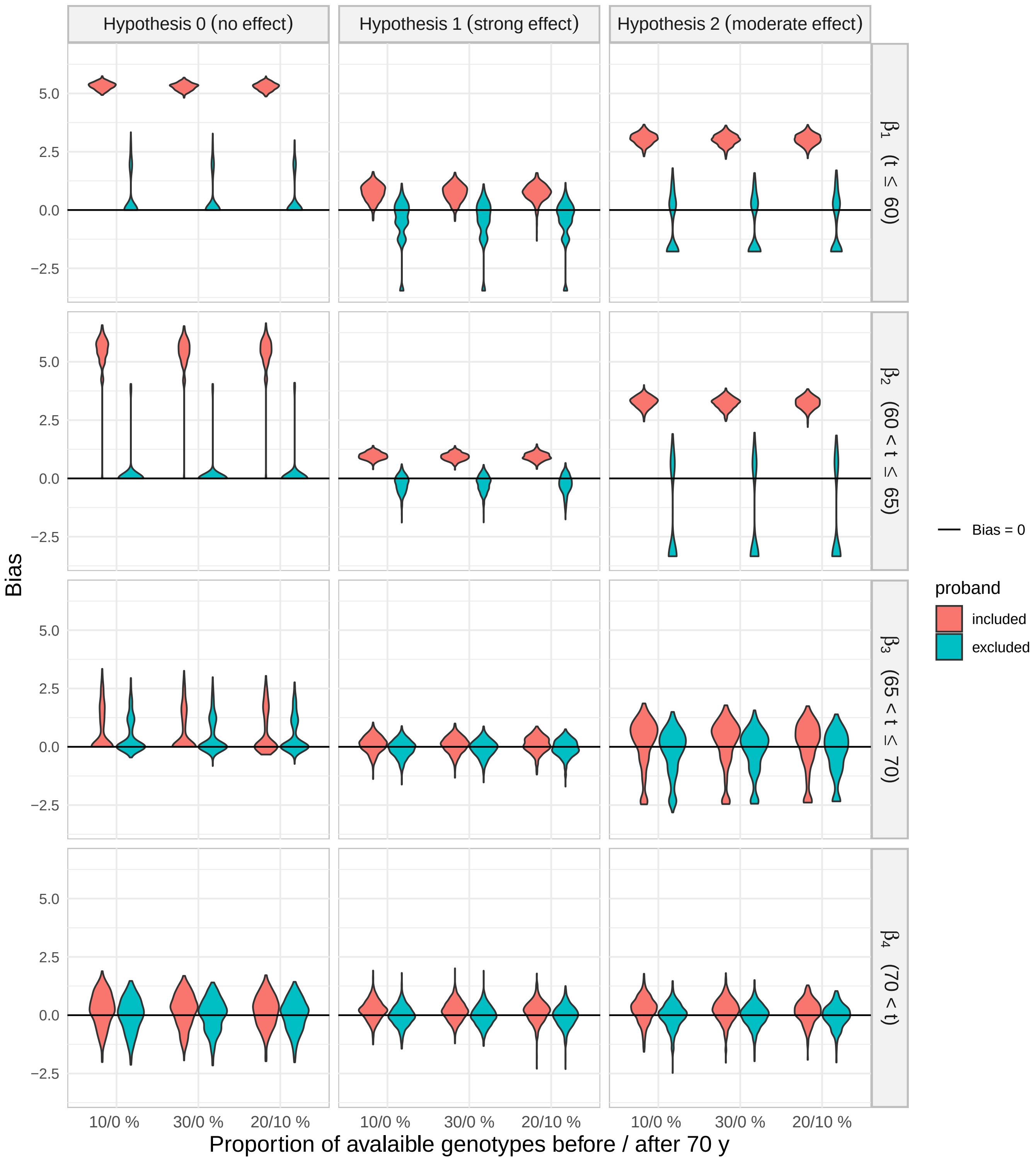


Figure S4: Results of simulation in scenario A when missingness depends on age

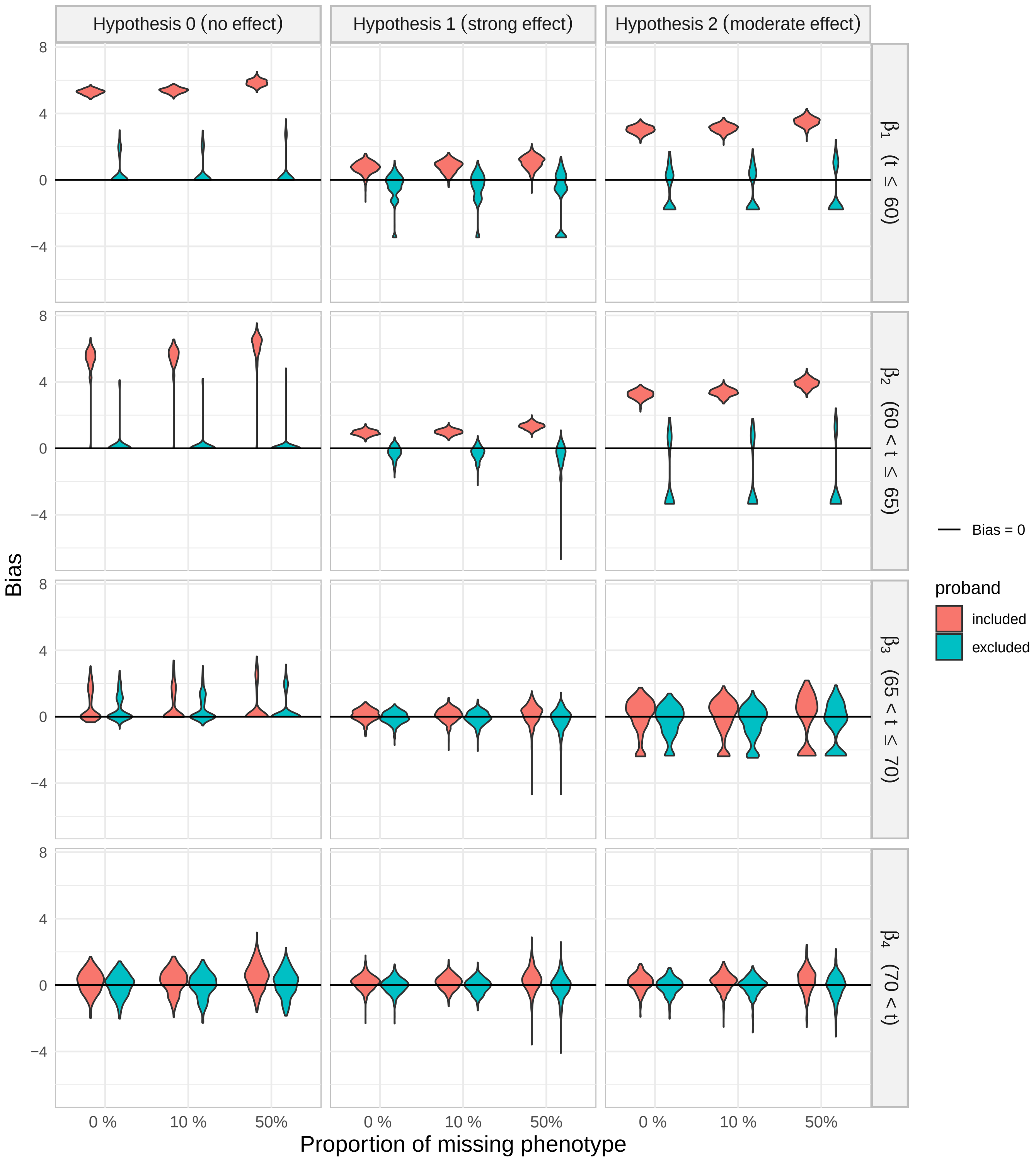


Figure S5: Results of simulation in scenario B

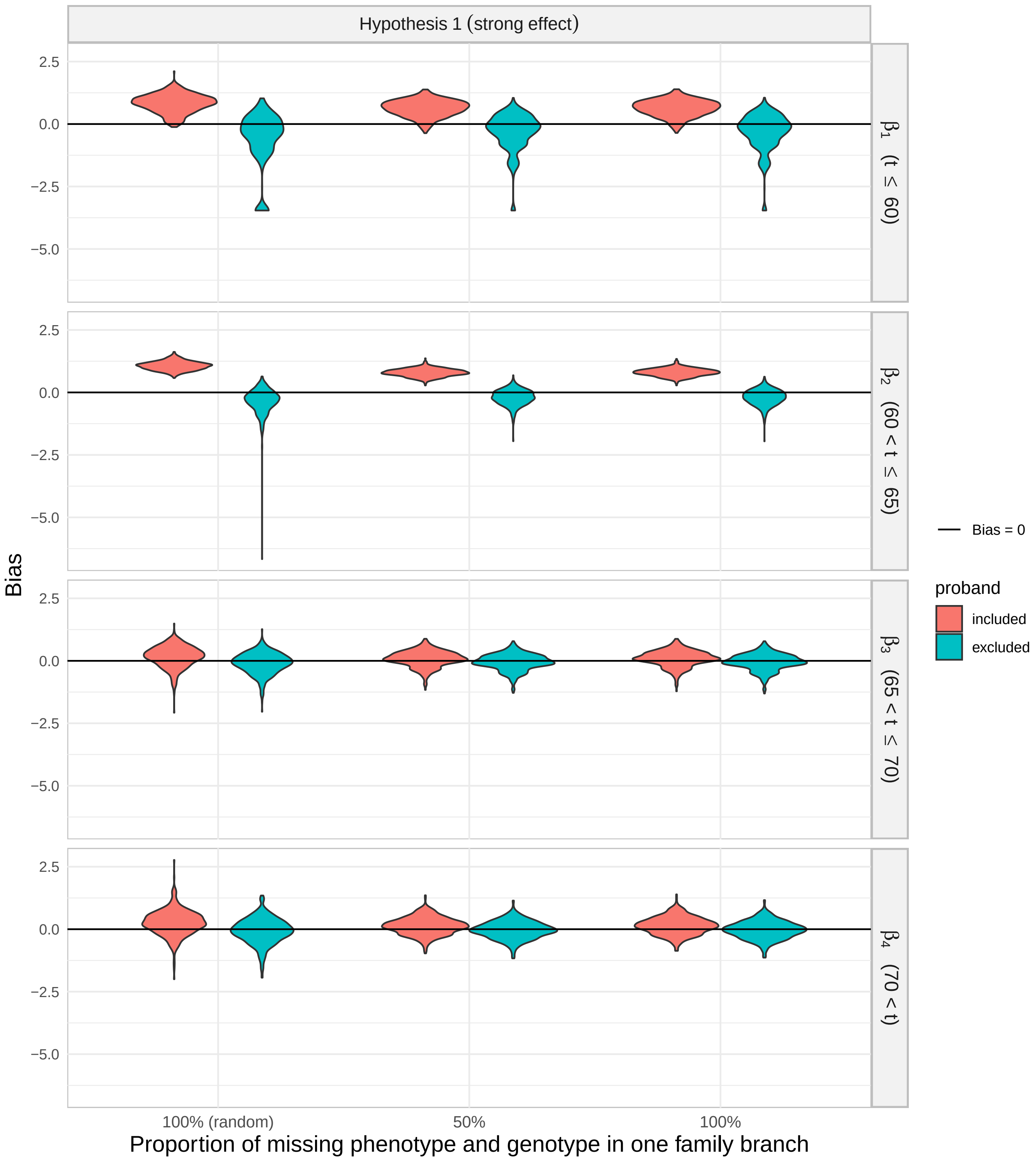


Figure S6: Results of simulation in scenario B when missingness is related to one family branch only (the one with the lower number of cases)

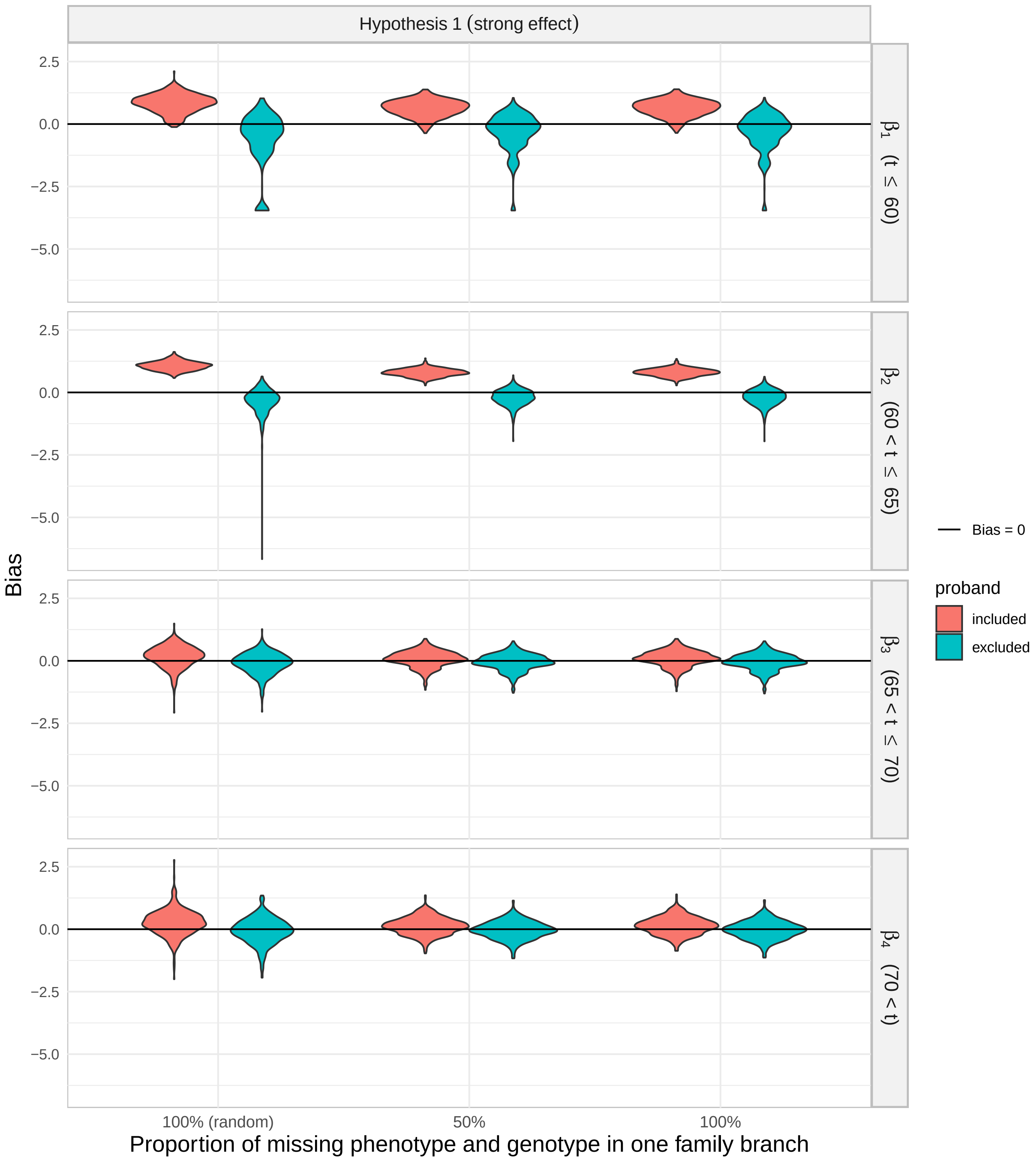


Figure S7: Results of simulation in scenario B when missingness is related to one family branch only (the one with parent not being a case)

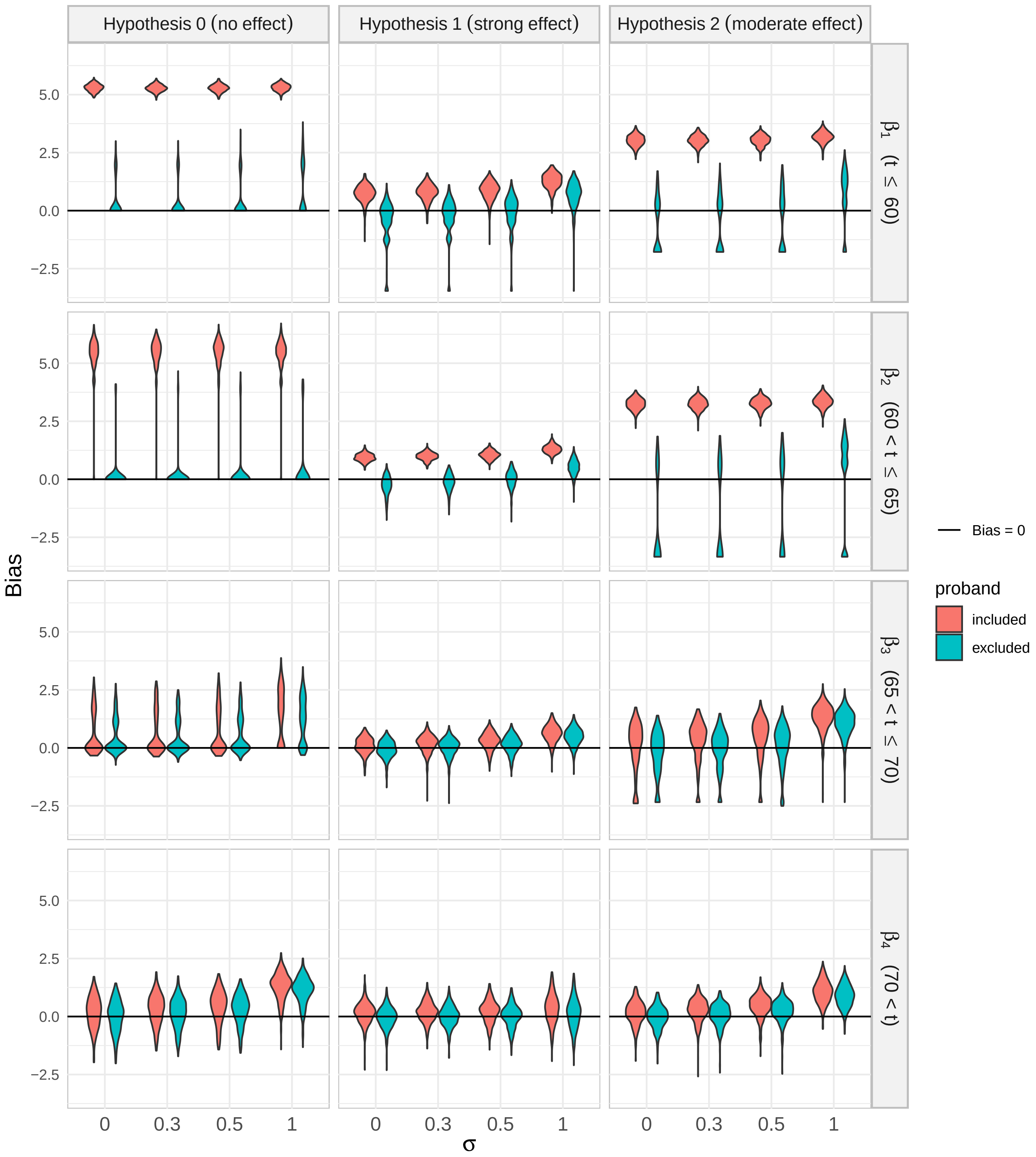


Figure S8: Results of simulation in scenario C with normal random effect

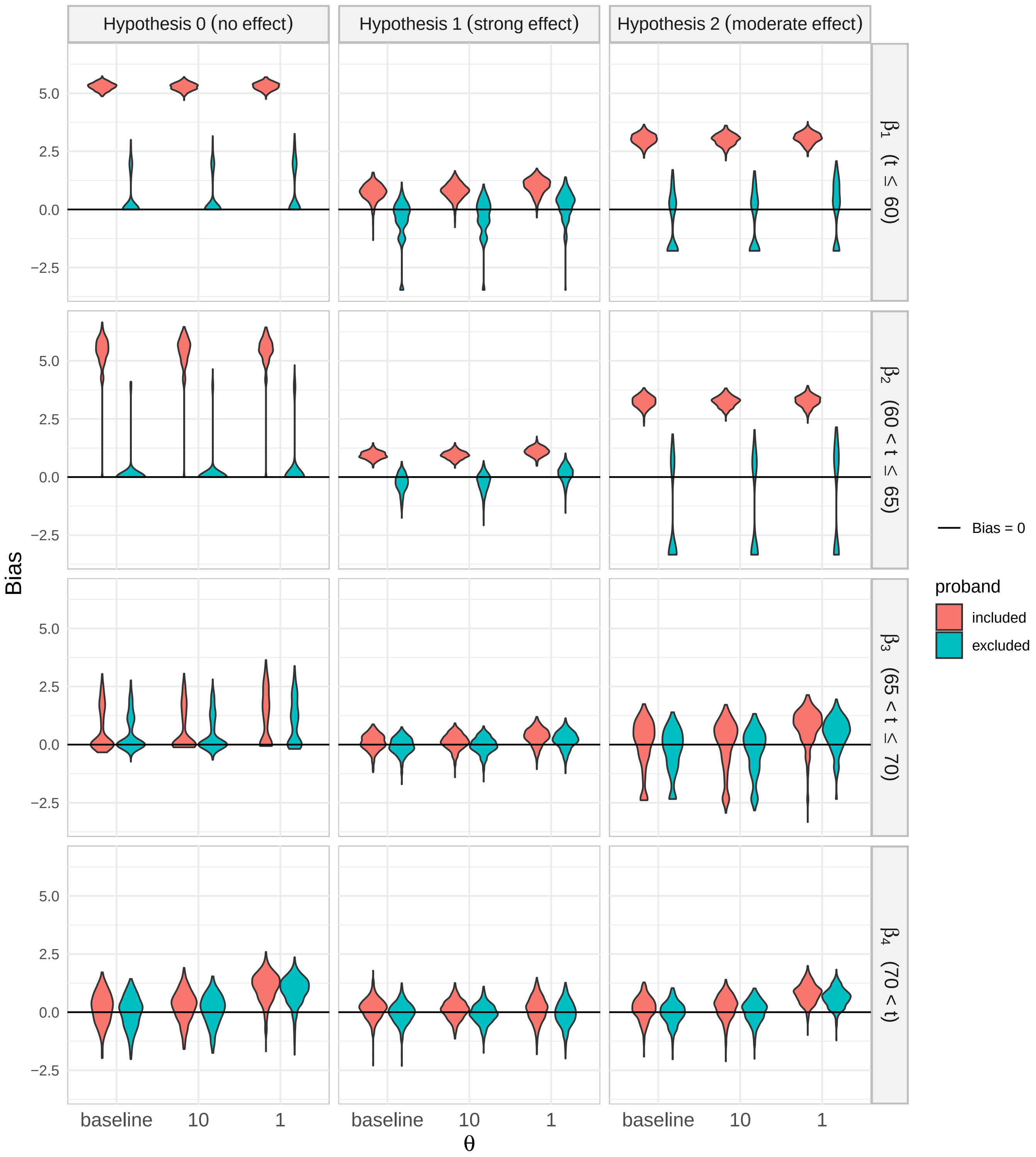


Figure S9: Results of simulation in scenario C with gamma random effect

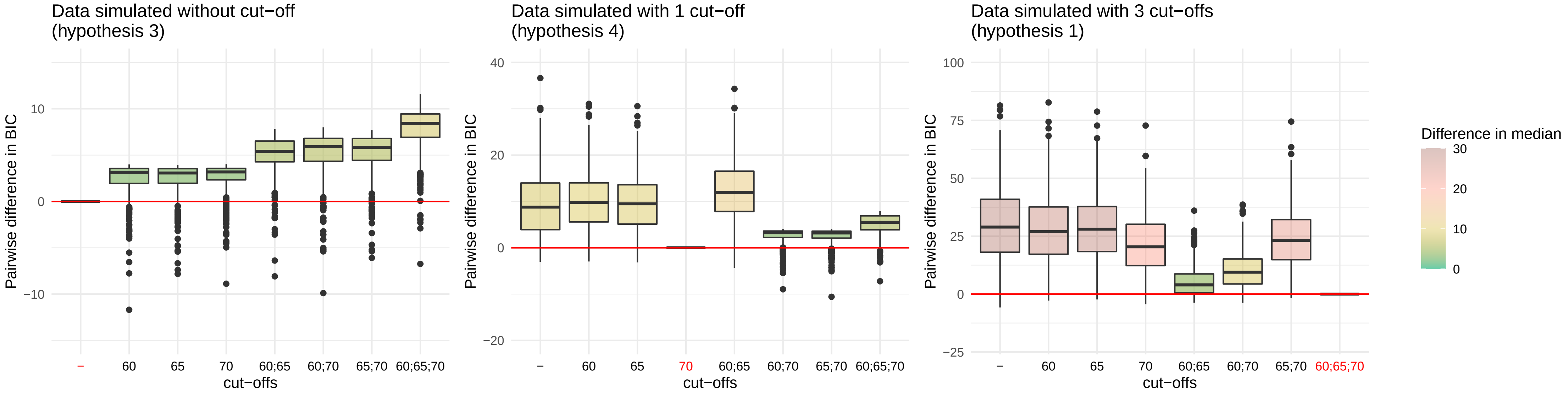


Figure S10: Results of simulation in scenario D

Y-axis represents the pairwise difference in BIC between the one obtained from the model using cut-offs displayed on the X-axix and the one obtained from the true cut-off (in red). To better compare boxplot within each graphic, Y-axis were not displayed on the same scales.

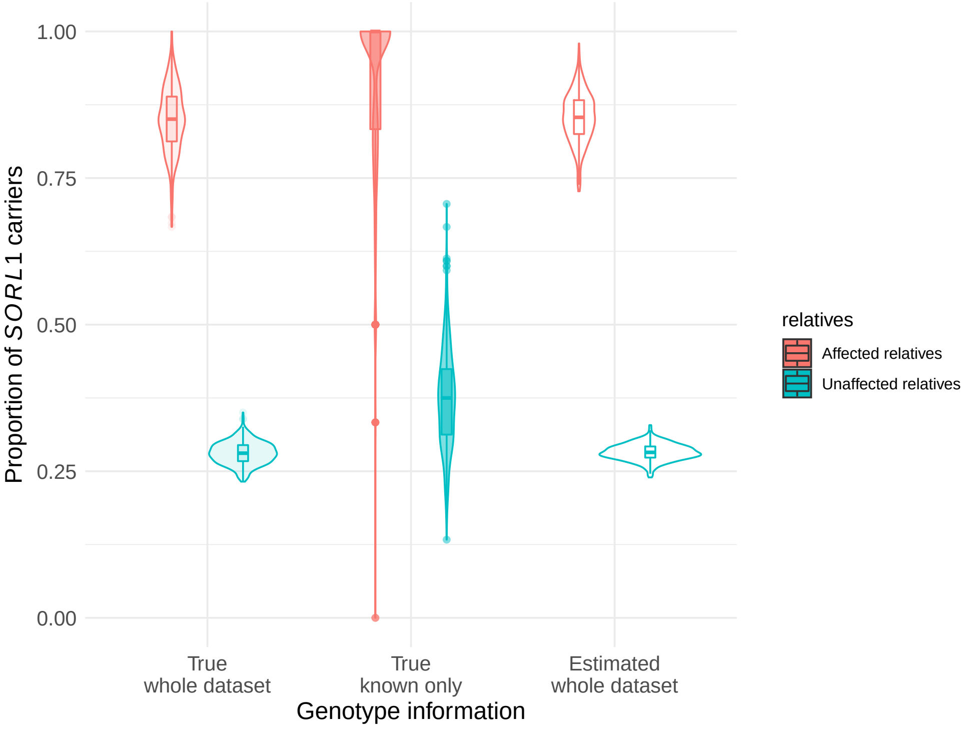


Figure S11: Posterior probability of carrying a variant in the baseline *scenario* over 500 simulated datasets. This graphic represents the proportion of carriers of *SORL1* variant of interest according to genotype information. ”True” refers to simulated genotypes and ”estimated” refers to the sum of posterior weights obtained in the E-step of our algorithm.

### 4 Supplemental Tables

| Type of variant | | Coding sequence | Protein sequence | Families |
|---|---|---|---|---|
| PTV | nonsense | c.802C>T | p.R268* | ROU-0699 |
| | | c.2412G>A | p.W804* | EFA-0067 |
| | | | | EFA-0280 |
| | | c.2473C>T | p.Q823* | EXT-1893 |
| | | c.2596C>T | p.R866* | EXT-1906 |
| | | c.2795G>A | p.W932* | ROU-1766 |
| | | c.3619C>T | p.R1207* | EXT-1231 |
| | | c.3647G>A | p.W1216* | ROU-1409 |
| | | c.3805C>T | p.Q1269* | SAL-0621 |
| | | c.4434C>A | p.C1478* | EXT-0050 |
| | | c.4963C>T | p.R1655* | EXT-1106 |
| | | c.5395C>T | p.R1799* | EXT-1756 |
| | | c.5463G>A | p.W1821* | EXT-0017 |
| | | c.6163G>T | p.E2055* | EXT-0087 |
| | | c.6279C>G | p.Y2093* | EFA-0198 |
| | frameshift | c.164delC | p.P55fs | EXT-0400 |
| | | c.1938delA | p.H647fs | ALZ-0167 |
| | | c.2603delC | p.T868fs | EXT-0438 |
| | | c.2882dupC | p.H962fs | EFA-0085 |
| | splice | c.1211+2T>G | | ROU-0055[a] |
| | | c.3947-3insG | | ROU-0055[a] |
| | | c.4213+1G>A | | RFA-0033 |
| | | c.4519+1G>A | | EXT-1023 |
| Mis3-LoF | | c.994C>T | p.R332W | EXT-0290 |
| | | c.1531G>C | p.G511R | EXT-0049 |
| | | c.1960C>T | p.R654W | ROU-0309 |
| | | | | ROU-1622 |
| | | | | ROU-1376 |

Table S1: *SORL1* variants included in our family cohort
PTV: protein-truncating variant; Mis3: Misense variant predicted as damaging by 3/3 software; LoF: loss-of-function; Mis3-LoF: Mis3 variant with *in vitro* LoF effect; a) Proband of ROU-0055 family is compound heterozygous for two distinct LoF variants (see Le Guennec et al.[2])

| Cut-offs | | | | BIC | |
|---|---|---|---|---|---|
| | | | | No censoring | Censoring |
| 60 | | | | 1173.41 | 1147.49 |
| | 65 | | | 1165.66 | 1148.52 |
| | | 70 | | 1155.37 | 1142.71 |
| | | | 75 | 1158.82 | 1149.04 |
| 60 | 65 | | | 1130.78 | 1113.25 |
| 60 | | 70 | | 1142.85 | 1129.81 |
| 60 | | | 75 | 1155.43 | 1145.47 |
| | 65 | 70 | | 1159.26 | 1146.64 |
| | 65 | | 75 | 1159.32 | 1149.70 |
| | | 70 | 75 | 1155.54 | 1145.98 |
| 60 | 65 | 70 | | 1124.24 | 1111.41 |
| 60 | 65 | | 75 | 1124.30 | 1114.55 |
| 60 | | 70 | 75 | 1143.23 | 1133.40 |
| | 65 | 70 | 75 | 1159.38 | 1149.85 |
| 60 | 65 | 70 | 75 | 1124.39 | 1114.71 |

Table S2: Comparison of models based on the Bayesian Information Criterion (BIC). Each row represents a model. Models differ in cut-offs for time interval definition in the piecewise constant function $\beta(t)$. No additional censoring was envisaged for our dataset. However, since only few individuals' observation time were greater that 85 years of age, we also compared BIC when data were censored at 85 years-old. Lower BIC means better fit. The same cut-offs were selected whether the data were censored or not censored (highlighted row).

| | Affected N = 61 | | Unaffected N = 246 | |
|---|---|---|---|---|
| *SORL1* carriers, $\mathbb{E}(N)$(%) | 48.60 (79.67%) | | 60.99 (24.79%) | |
| *APOE* × *SORL1* (+ vs WT) | + | WT | + | WT |
| ε4 non-carriers, $\mathbb{E}(N)$ (%) | 20.87 (34.21%) | 5.49 (9.00%) | 39.26 (15.96%) | 115.04 (46.76%) |
| ε4 heterozygous carriers, $\mathbb{E}(N)$ (%) | 22.61 (37.07%) | 5.79 (9.49%) | 18.70 (7.60%) | 63.30 (25.73%) |
| ε4ε4 carriers, $\mathbb{E}(N)$ (%) | 5.12 (8.39%) | 1.13 (1.85%) | 3.03 (1.23%) | 6.67 (2.71%) |

Table S3: Expected total number of individual by genotype.
The number of expected carriers $\mathbb{E}(N)$ of each genotype was estimated by summing posterior weights obtained in the E-step of our algorithm. Results are given for individuals with informative phenotype (i.e. status and age $\geq$ 40 years-old).
